# Characterization of a vascular bundle localizing *Gossypium hirsutum* NAC4 transcription factor promoter for its role in environmental stress responses

**DOI:** 10.1101/510578

**Authors:** Vikas Shalibhadra Trishla, Sureshbabu Marriboina, Prasanna Boyidi, Padmaja Gudipalli, Pulugurtha Bharadwaja Kirti

**Author notes:** Corresponding author email id. Faculty of Basic Sciences and Humanities, Rajendra Prasad Central Agricultural University, Pusa - Samsthipur, Bihar, India and Agri Biotech Foundation, Rajendranagar, Hyderabad, India.

## Abstract

We have studied the expression of *GhNAC4*, a NAC domain-containing transcription factor from *Gossypium hirsutum*. The expression of *GhNAC4* was upregulated by ABA, cytokinin, JA, GA, auxin, and ethylene. Its expression was also highly induced by drought, osmotic stress, oxidative stress, salinity, high and low-temperature stress, and wounding. To corroborate these observations, we isolated the promoter of *GhNAC4* and fused it transcriptionally with *uidA* (GUS) gene for analyzing its patterns of expression in transgenic tobacco. The Promoter-GUS fusion was also induced by various phytohormones and environmental stresses. The spatio-temporal analysis of the promoter of the *GhNAC4* gene revealed that GUS expression was mostly localized to the vascular bundles along with shoot apical meristem and guard cells. We also observed intense staining in other cells upon wounding. A sequence analysis of the promoter revealed the presence of several motifs pertaining to phytohormone responsiveness, stress-inducibility, light and sugar-responsiveness and tissue-specificity. These data were corroborated by a detailed bioinformatic analysis of the promoter sequence of *GhNAC4* for identifying the conserved sequences that are associated with the expression of genes in a spatio-temporal or inducive manner. All these data suggests that GhNAC4 is a vascular tissue localizing NAC transcription factor, which might act as a node integrating environmental stress signals for modulating plant growth and development with the aid of phytohormonal stimuli.

**Key message:** GhNAC4 transcription factor from cotton localizes to vascular bundles and is highly upregulated by phytohormones and environmental stresses.

## Introduction

In plants, vascular tissues consist of xylem and phloem which are derived from the meristematic vascular procambium and cambium (Esau 1977). The proliferation of cambium, development, differentiation, and patterning of vascular tissue requires hormonal responses, transcriptional regulators, and peptide signalling components (Růžička et al. 2015). Various transcription factors (TFs) like NAC, MYB, and HD-ZIP are implicated in vascular bundle development (Yang and Wang 2016).

NAC (NAM ATAF CUC) TFs constitute one of the largest plant-specific TF super-families with 171 and 151 members in Arabidopsis and rice, respectively. (Nuruzzaman et al. 2010). So far 283 NAC genes have been identified in *Gossypium hirsutum*, 147 in *G. arboreum*, 267 in *G. barbadense*, and 149 in *G. raimondii*(Sun et al. 2018). A typical NAC TF carries two domains, a conserved N-terminal domain and a highly divergent C-terminal domain (Olsen et al. 2005). Previous studies have shown that NAC TFs play essential roles in regulating a wide variety of biological processes such as shoot apical meristem development (Souer et al. 1996), seed development (Sperotto et al. 2009), leaf senescence (Guo and Gan, 2006), flower development (Sablowski and Meyerowitz, 1998), fibre development (Ko et al. 2007), abiotic (Hu et al. 2006) and biotic stress responses (Nakashima et al. 2007).

The biological functions of NAC TFs in vascular development have been emerging recently. NAC TFs act as master switches regulating the vascular tissue differentiation and secondary cell wall formation (Wang and Dixon 2012). VASCULAR-RELATED NAC-DOMAIN1 (VND1) to VND7 are specifically expressed in vascular tissues. VND6 and VND7 regulate vessel differentiation at inner metaxylem and protoxylem in the roots respectively (Kubo et al. 2005). NAC SECONDARY WALL THICKENING PROMOTING FACTOR1 (NST1), NST2, and SECONDARY WALL-ASSOCIATED NAC DOMAIN PROTEIN 1 (SND1)/NST3/ANAC012 act as transcriptional switches that regulate secondary cell wall thickening in various tissues and are localized to xylary and extraxylary fibres in the inflorescence stem (Ko et al. 2007; Mitsuda et al. 2005; Mitsuda et al. 2007). XYLEM NAC DOMAIN 1 (XND1) is specifically expressed in xylem and negatively regulates tracheary element growth (Zhao et al. 2008).

So far, very little information is available on the role of NAC TFs in vascular development in cotton. GhXND1, a cotton NAC TF is involved in negative regulation of xylem development. Ectopic expression of GhXND1 resulted in a decrease in the number of xylem vessels and reduction of interfascicular fibre thickness (Li et al. 2014). GhFSN1 (fibre secondary cell wall-related NAC1) is expressed in cotton fibres and is vital for secondary cell wall synthesis (Zhang et al. 2018). However, progress toward related research has been slow. Comprehensive expression analysis of NAC TFs would be useful in identifying and understanding the key mechanistic steps in their biological function.

Promoters of only a few NAC TFs mainly from Arabidopsis and Rice, have been studied for their induction and localisation patterns. *OsNAC6* promoter-GUS fusion was induced by ABA, MeJA and various environmental stresses. And the promoter sequence contained several *cis*-acting elements involved in the response to abiotic and biotic stresses (Nakashima et al. 2007). *OsNAC5* promoter was induced by ABA and was localized to roots and leaves under ABA and NaCl treatments (Takasaki et al. 2010). The expression patterns of *SNAC1* promoter was observed in callus, root, leaf, guard cells, ligule, stamen, and pistil (Hu et al. 2006) while *SNAC2* was observed only in roots and internodes (Hu et al. 2008). The Promoter-GUS fusion of *NST1* exhibit localisation to the anthers, filaments of stamens, and carpels and vascular bundles of the leaf, while *NST2* was mostly localized to the anther wall and pollen grains (Mitsuda et al. 2005). The expression patterns of *ORE1*/*ANAC092*/*AtNAC2* (*oresara1*) and *ORS1* (ORESARA1 SISTER1) promoters was observed in cotyledons, senescent leaves roots and mature floral tissues (Balazadeh et al. 2010, 2011).

In the present study, cotton (*Gossypium hirsutum* L., belonging to the family, Malvaceae), which is one of the most widely cultivated and economically important crops was investigated. Cotton is a primary source of natural fibre and an important oilseed crop. However, cotton-growing areas are prone to high and low temperatures, extreme drought, salinity, and pest infestation that can hinder crop productivity (Ullah et al. 2017). Very little information is available on the molecular mechanisms that regulate the responses and adaptation of the cotton plant to environmental stresses. Hence, advancements to accelerate stress tolerance are very important for cotton production.

Based on the cDNA sequence of NAC4 from cotton (*GhNAC4*) cloned previously by Meng et al. (2009), we undertook the present study. As a significant step towards elucidating the mechanisms that regulate *GhNAC4* expression, the promoter region was cloned for the first time in our study, and the *cis*-acting elements have been predicted subsequently. To determine the participation of GhNAC4 in environmental stress responses via phytohormonal stimuli and to identify the key mechanistic steps for improving tolerance, we also examined the expression of *GhNAC4* and induction of *GhNAC4* promoter under various environmental and phytohormonal treatments. We studied the spatio-temporal localisation of GhNAC4 through Promoter-GUS fusion for the first time in a heterologous system, tobacco and found that it preferentially localizes to vascular bundles. To the best of our knowledge, GhNAC4 is one of the very few reported NAC TFs that are expressed in vascular bundles in cotton. All the observations presented here would allow us to gain more insight into the molecular function and regulation of NAC transcription factors in cotton. Such a study would also provide valuable information for future breeding projects to enhance cotton yield under stress conditions.

## Materials and methods

### Plant material and stress treatments

Cotton (*Gossypium hirsutum* var. JK Durga) seeds were surface sterilized with 70% ethanol for 2 min followed by 4% sodium hypochlorite for 15 min. The seeds were then rinsed four times with sterilized water and soaked for 5-6 h. Subsequently, they were germinated and grown for two weeks on sterile blotting paper placed on top of 0.5X MS media without any growth regulators.

For hormonal treatments, the filter paper on which the seedlings were grown, was moistened with either 100 µM ABA, 100 µM MeJA, 100 µM SA, 20 µM 6-BAP, 20 µM GA_3_ or 20 µM IAA and incubated for 24 h prior to sampling. As controls, untreated seedlings and filter paper moistened with water having the same concentration of sodium hydroxide or alcohol used for dissolving the hormones were used. Ethylene treatment was carried out for 24 h by placing the seedlings on filter paper in a sealed container, and Ethephon was added to the box and diluted to a final concentration of 10 ppm in distilled water. Seedlings in a similar container having air were used as a control.

High salt and osmotic stress were induced by moistening the filter paper with 0.3 M NaCl and 0.3 M mannitol respectively, and the seedlings were allowed to grow for 24 h. Oxidative stress was induced by moistening the filter paper with 10 µM methyl viologen and incubated for 24 h. For inducing drought stress, the seedlings were incubated on filter paper moistened with 15% (w/v) PEG 6000 for 24 h. Air drying stress was carried out by placing the seedlings on the surface of dry filter paper for 30 min. Flooding stress was achieved by immersing the seedlings in distilled water for 24 h. For the wounding treatment, leaves of the seedlings were squeezed with a forceps, and the wounded leaves were harvested after 30 min. For the combination of dark and cold treatments, the seedlings were wrapped in aluminium foil and incubated at 4 °C for 24 h, while the dark treatment was carried out at 25 °C for 24 h. In the high-temperature stress, the seedlings were subjected to 42 °C for 12 h. Following all the treatments, the leaves were quickly frozen in liquid nitrogen to analyse the expression levels of *GhNAC4* gene.

To determine the degree of promoter activation, homozygous T_2_ tobacco seedlings were subjected to different hormonal treatments and environmental stresses following the same methodology as applied to cotton seedlings, for GUS activity measurement.

### RNA isolation and real-time quantitative PCR

Total RNA was extracted from the leaves of control and treated cotton seedlings by a CTAB (cetyltriethylammonium bromide) extraction procedure as described by Chang et al. (1993). To avoid DNA contamination, the total RNA was treated with RNase free DNase1 (Epicentre Biotechnologies, USA) by incubating at 37 °C for 15 min. One µg of total RNA was used for synthesising the first-strand cDNA using RevertAid 1^st^ strand cDNA synthesis kit (Thermo Fischer Scientific, USA) following the manufacturer’s instructions. An oligo-dT_18_ was used as the primer in the reverse transcription reaction. For the real-time quantitative PCR, cDNA was diluted to 100 ng/µl and was mixed with the SYBR master mix (Kapa Biosystems, USA), and amplification was carried out according to the manufacturer’s protocol in the StepOne Plus machine (Applied Biosystems, USA). The constitutively expressing Ubiquitin gene (*GhUBQ7*, GenBank accession no. DQ116441) from cotton, was used as an internal reference gene. The primer sequences used to amplify the internal regions of *GhNAC4* (NAC4-RTF and NAC4-RTR), and *GhUBQ7* (UBQ7-RTF and UBQ7-RTR, Kuppu et al. 2013) are mentioned in Table 1. To ensure the gene specificity, the amplicons obtained by these primers were confirmed by sequencing. The fold change in the *GhNAC4* gene levels relative to *GhUBQ7* gene was determined using the ΔΔC_T_ method (Livak and Schmittgen 2001). The experiment was performed in triplicates, and two independent biological replicates were used in the analysis.

**Table 1.**
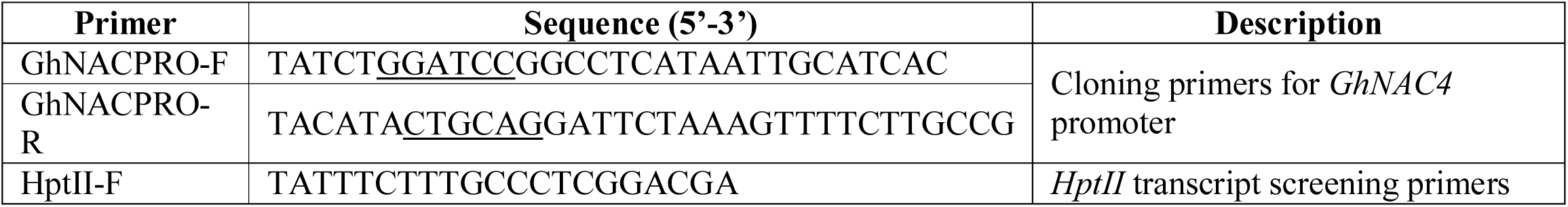
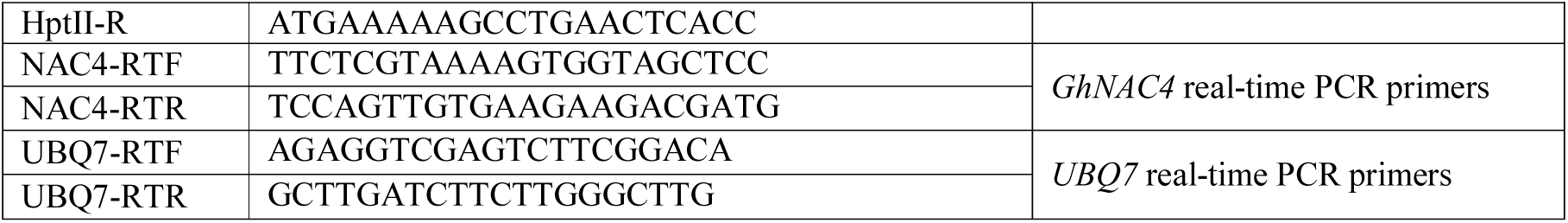
Primer sequences used in the experiments. Underlined sequences are restriction enzyme recognition site

### Isolation and cloning of GhNAC4 promoter from cotton

The full-length CDS of *GhNAC4* (GenBank accession number EU706342.1) was used as a query to retrieve the 5’ upstream sequence from Phytozome (https://phytozome.jgi.doe.gov/). Genomic DNA was isolated from the leaves of cotton by the CTAB method (Murray and Thompson 1980). It was used as a template for PCR amplification of a DNA fragment, in the range of −1492 bp to +119 bp (relative to the transcription start site). The primer sequences (GhNACPRO-F, GhNACPRO-R) used for the amplification are mentioned in Table 1. The amplicon was cloned into pTZ57R/T (ThermoFisher Scientific, USA) and the accuracy was verified by sequencing. The fragment was excised and sub-cloned into *Bam*HI and *Pst*I restriction sites of the promoter-less vector, pCAMBIA 1381Z (Cambia, Australia) to generate a transcriptional fusion comprising the 5’ upstream region of *GhNAC4* and *uidA* gene (pPRO*_GhNAC4_*:GUS). This construct was used for *Agrobacterium tumefaciens* (strain EHA105) transformation by using the freeze-thaw method.

### Generation of tobacco transgenics

*In vitro* grown three weeks old *Nicotiana tobacum* cv. Samsun leaves were used as explants for transformation with Agrobacterium harbouring either the empty pCAMBIA 1381Z (vector control) or the pPro*_GhNAC4_*:GUS vector, following to Horsch et al. (1985). Explants were co-cultivated with Agrobacterium on a co-cultivation medium (MS salts with 2 mg/l BAP, 0.1 mg/l NAA and 3% sucrose pH 5.8) for 48 h in dark and later were transferred to the shoot induction medium (MS salts with 2 mg/l BAP, 0.1 mg/l NAA, 3% sucrose, 10 mg/l hygromycin and 250 mg/l Cefotaxime pH 5.8) with 16/8 h light/dark photoperiod at 24±2 °C. The regenerated shoots were sub-cultured on shoot elongation medium (MS salts with 1 mg/l BAP, 0.05 mg/l NAA, 3% sucrose, 15 mg/l hygromycin and 250 mg/l Cefotaxime pH 5.8). The hygromycin resistant shoots were further sub-cultured on rooting medium (0.5xMS salts with 0.1 mg/l NAA and 20 mg/l hygromycin pH 5.8). Plants with well-developed roots were transferred to sterile soil in small plastic cups for acclimatisation.

Further, well-established plants were shifted to pots and allowed to grow to maturity in a green house and set seeds. The putative primary tobacco transformants (T_0_) were screened by genomic PCR for the presence of *GhNAC4* promoter sequence and hygromycin gene (*HptII*) (with primers, HptII-F and HptII-R; see Table 1) and allowed to self-pollinate and set seeds. The T_1_ seeds from selected plants were germinated on 0.5X MS medium supplemented with 25 mg/l hygromycin. Copy number of the DNA integration was determined by Hyg^R^ segregation test to select the single copy integration events that give a ratio similar to the monohybrid ratio of resistant and susceptible seedlings. Segregation analysis was carried out by counting the number of green and bleached seedlings. The hygromycin-resistant (green) seedlings were later transferred to soil in the green house after 2-3 weeks and allowed to set seeds. The T_2_ progeny were germinated on 0.5X MS medium supplemented with 25 mg/l hygromycin to identify the homozygous single copy integration T_2_ lines.

### Histochemical localisation and fluorometric measurement of GUS activity

β-glucuronidase (GUS) activity was assayed as described by Jefferson et al. (1989) with minor modifications. For histochemical staining, the tissues were vacuum infiltrated with the solution containing 1 mM X-Gluc (5-bromo-4-chloro-3-indolyl-b-D-glucuronide), 1 mM potassium ferrocyanide, 1 mM potassium ferricyanide, 1 mM EDTA and 0.1% Triton X-100 in 50 mM phosphate buffer (pH 7.0) and were incubated at 37 °C for 12-14 h in the dark. After staining, the tissues were fixed in a solution containing 4% formaldehyde in 50 mM phosphate buffer (pH 7.0) for 12 h at 4 °C and subsequently cleared in 70% ethanol at room temperature. Photographs were taken using M165 FC and DM6B microscopes (Leica Microsystems, Germany).

For the fluorometric assay, the tissue was homogenised in 400 µl GUS extraction buffer containing 10 mM EDTA and 0.1% Triton X-100 in 50 mM phosphate buffer (pH 7.0). After centrifugation at 12,000 rpm (4 °C) for 15 min, 5 µl of homogenate was diluted with 95 µl of extraction buffer and mixed with 100 µl of extraction buffer having 2 mM 4-methyl-umbelliferyl-β-D-glucuronide (4-MUG, Duchefa, Netherlands) and incubated at 37 °C for 1 h. The reaction was terminated by the addition of 1.8 ml of 200 mM sodium carbonate. Total protein concentration in the homogenate was assessed by the Bradford method (Bradford, 1976) with Bovine Serum Albumin (BSA) as a standard. Fluorescence (excitation 363 nm, emission 447 nm) was determined by Infinite 200 plate reader (Tecan, Switzerland) and GUS activity was expressed as pmol of 4- methyl-umbelliferone (4-MU, Sigma, USA) per µg protein per min. 4-MU in the range of 20 nM-100 µM was used to generate a standard curve. Each MUG assay was performed in triplicate and repeated three times.

### Bioinformatics analysis of *GhNAC4* promoter sequence

A search for the putative *cis*-acting regulatory elements in the putative promoter sequence was conducted using the PlantPAN 2.0 (Chang et al. 2008, http://plantpan2.itps.ncku.edu.tw/), PLACE (Higo et al. 1999, http://www.dna.affrc.go.jp/htdocs/PLACE/) and PlantCARE (Lescot et al. 2002, http://bioinformatics.psb.ugent.be/webtools/plantcare/html/) databases.

### Statistical analysis

All experiments were repeated at least three times, and the data were expressed as the mean ± SE. Error bars shown are the standard error of the experimental data. Data were- analysed by one-way analysis of variance (ANOVA) using SigmaPlot 11.0 software. *** P<0.001, ** P□<□0.01 and * P□<□0.05 represents significant differences at 0.1, 1 and 5% level respectively. ‘ns’ represents no significant difference.

## Results

### Expression analysis revealed the responsiveness of *GhNAC4* gene to various phytohormones and environmental stresses

Plant stress responses are thought to be regulated by various hormones and multiple signalling pathways. They show significant overlap with the gene expression patterns (Singh et al. 2002). To gain an insight into the impact of environmental stresses and phytohormones on gene expression of *GhNAC4*, real-time expression analysis was carried out by subjecting cotton seedlings to various treatments (see materials and methods). Fig. 1 shows that *GhNAC4* responded differentially to treatments with several phytohormones. Phytohormones like BAP, ABA, GA3, and MeJA significantly upregulated the expression of *GhNAC4*. BAP caused the induction of expression by 10.4 fold, GA3 by 11.5 fold, ABA by 6.9 fold and MeJA by 7.8 fold at 24 h. However, other hormones like IAA, Ethylene and SA caused only a weaker up-regulation of its expression (4-5 fold) at 24 h.

**Fig. 1.**
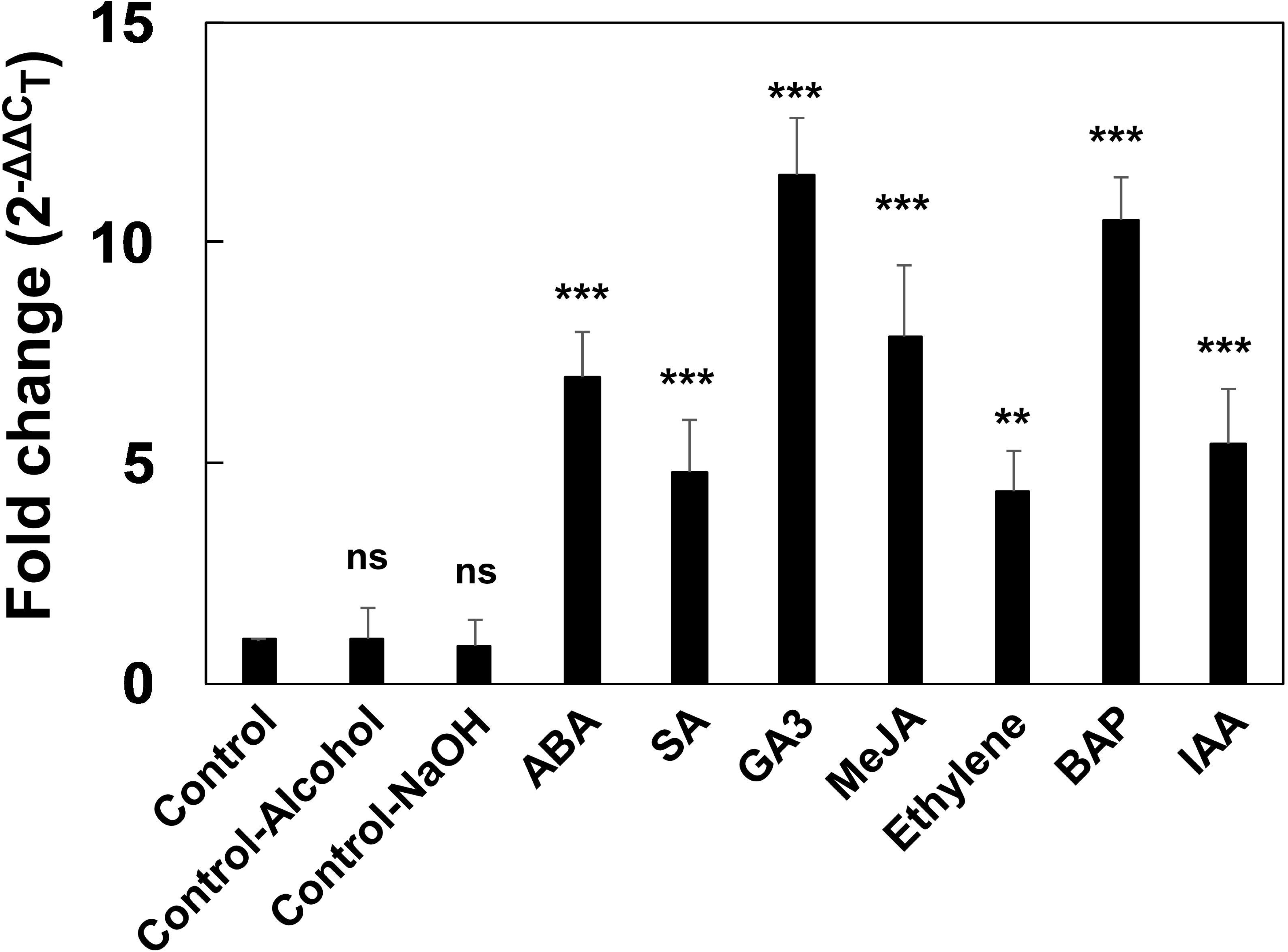
Expression patterns of *GhNAC4* transcript in response to various phytohormones. qRT-PCR expression analysis of the *GhNAC4* gene in *G. hirsutum* leaves after ABA, MeJA, SA, 6-BAP, GA_3_, IAA, or Ethephon treatment. Two weeks-old cotton seedlings incubated for 24 h under the treatment, were used for analysis. mRNA levels of GhNAC4 gene were normalized to that of Ubiquitin gene, *GhUBQ7*. The data are shown as the means ± SE (n=3). A statistical analysis with one-way ANOVA indicates significant differences (** P<0.01, *** P<0.001, ns - not significant)

PEG-induced drought treatment resulted in very high up-regulation of *GhNAC4* expression (∼184 fold) at 24 h. Other abiotic stress treatments like high salinity and osmotic stress (caused by mannitol) also led to high up-regulation of expression (43.6 and 58.7 fold respectively). Air-drying caused 10.2 fold, cold 19.4 and MV by 14.5 fold up-regulation in the *GhNAC4* transcripts. However, *GhNAC4* was weakly upregulated by high temperatures, flooding, wounding and dark treatments (2-6 fold) compared to other treatments as shown in Fig. 2.

**Fig. 2.**
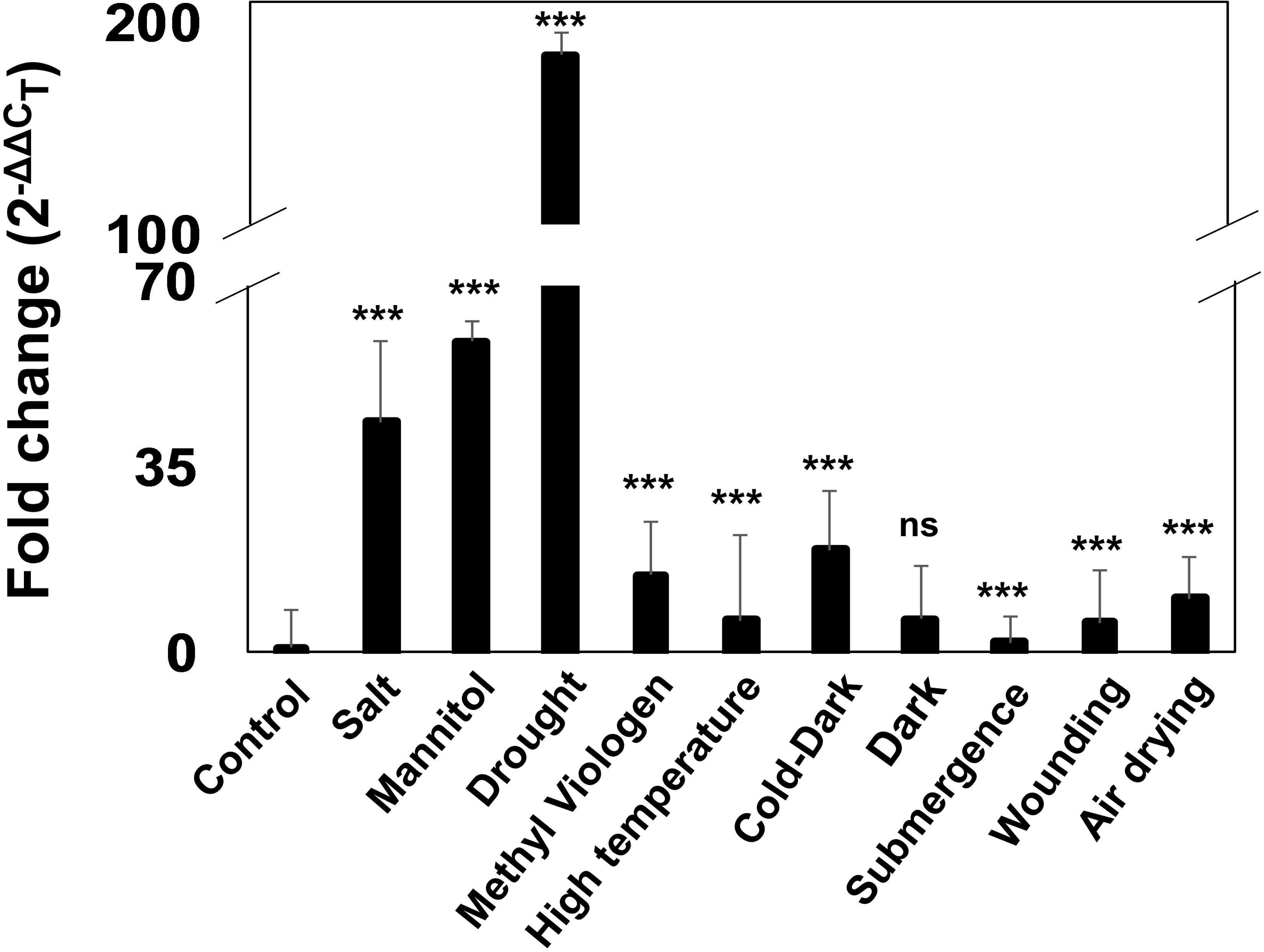
Expression patterns of *GhNAC4* transcript in response to various environmental stresses. qRT-PCR expression analysis of the *GhNAC4* gene in *G. hirsutum* leaves after NaCl, mannitol, PEG, methyl viologen, high temperature, air drying, submergence, wounding, dark or combination of dark and cold treatment. Two weeks-old cotton seedlings were used for analysis. mRNA levels of *GhNAC4* gene were normalized to that of Ubiquitin gene, *GhUBQ7*. The data are shown as the means ± SE (n=3). A statistical analysis with one-way ANOVA indicates significant differences (*** P<0.001, ns - not significant)

### Sequence analysis of *GhNAC4* promoter

A DNA fragment of 1612 bp corresponding to the upstream regulatory region (−1492 bp to +119 bp)of the *GhNAC4* was amplified from *G. hirsutum* (Var. JK Durga) and was confirmed by sequencing. Sequence similarity of 91.6% (LALIGN, https://www.ebi.ac.uk/Tools/psa/lalign/) was observed between the promoter region of *G. raimondii* and *G. hirsutum* of *NAC4* genes. No significant difference was observed between the predicted motifs in the two sequences. The composition of the GC content of *GhNAC4* promoter was 29.9% (BioEdit, Hall 1999), which is in accordance with the observed range (Joshi 1987) for a plant promoter. The putative transcription start site (TSS) predicted by Softberry database (www.softberry.com) using the default settings was located 119 bp upstream of the ATG translation start codon, which was consistent with the features of a eukaryotic promoter as shown in Fig. 3. The predicted TATA box was located 16 bp upstream of TSS, and a CAAT box was located 179 bp upstream of TATA box. Several other CAAT boxes were also predicted in the entire length of the sequence.

**Fig. 3.**
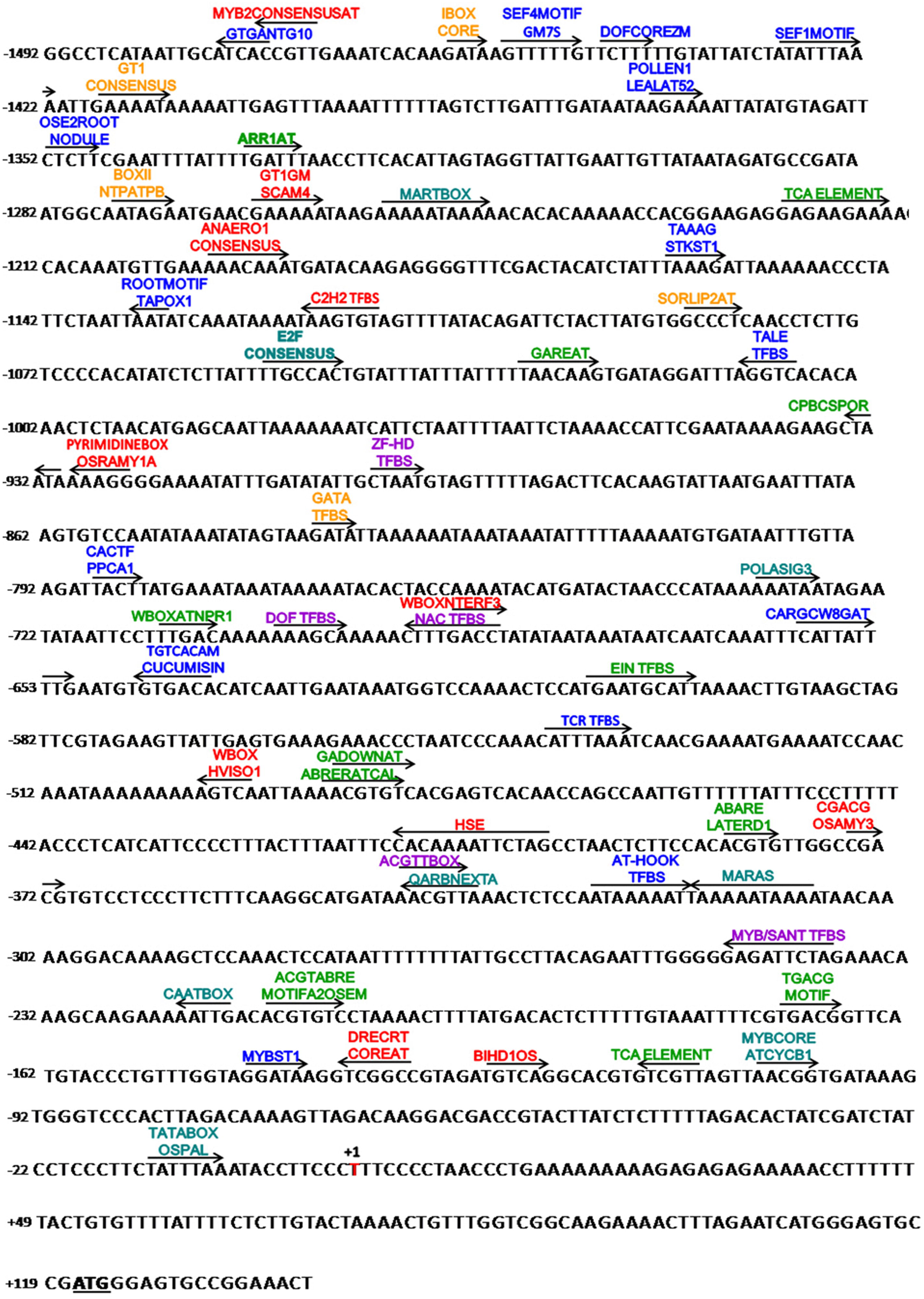
Physical map of *GhNAC4* promoter. Nucleotide sequence of the 1492 bp upstream region of *GhNAC4* gene from *G. hirsutum* var JK Durga. The putative transcription start site is in bold and is designated as +1. The translation start site is bolded and underlined. The numbers on the left side indicate the distance from the transcription start site. The sequence was analysed by PLACE, PlantCARE and PlantPAN2.0 programs. All the predicted motifs are indicted by arrow and their names are mentioned above. ‘→’ and ‘←’ indicates that the predicted motif is on positive (5’-3′) and negative (3′-5’) strand respectively. Stress inducible motifs are represented in red, phytohormone responsive motifs are in green, light inducible motifs are in orange, tissue specificity motifs are in blue, transcriptional related motifs in purple and transcription factors binding sites are in teal colour

### Multiple *cis*-acting elements were predicted on the *GhNAC4* promoter

Putative *cis*-acting regulatory elements and their location were searched using the PlantPAN 2.0, PlantCare and PLACE software tools. Only statistically significant motifs (P value > 0.9) were selected. The resulting putative *cis*-acting regulatory elements were grouped into six classes, as shown in Table 2 including phytohormone-responsive motifs, stress-responsive motifs, light responsive, basic transcription elements, tissue-specific elements, and other TF binding sites. We observed a relatively high proportion of tissue-specific motifs and transcription factors binding sites followed by light specific, hormone responsive and stress responsive elements (Fig. 4).

**Fig. 4.**
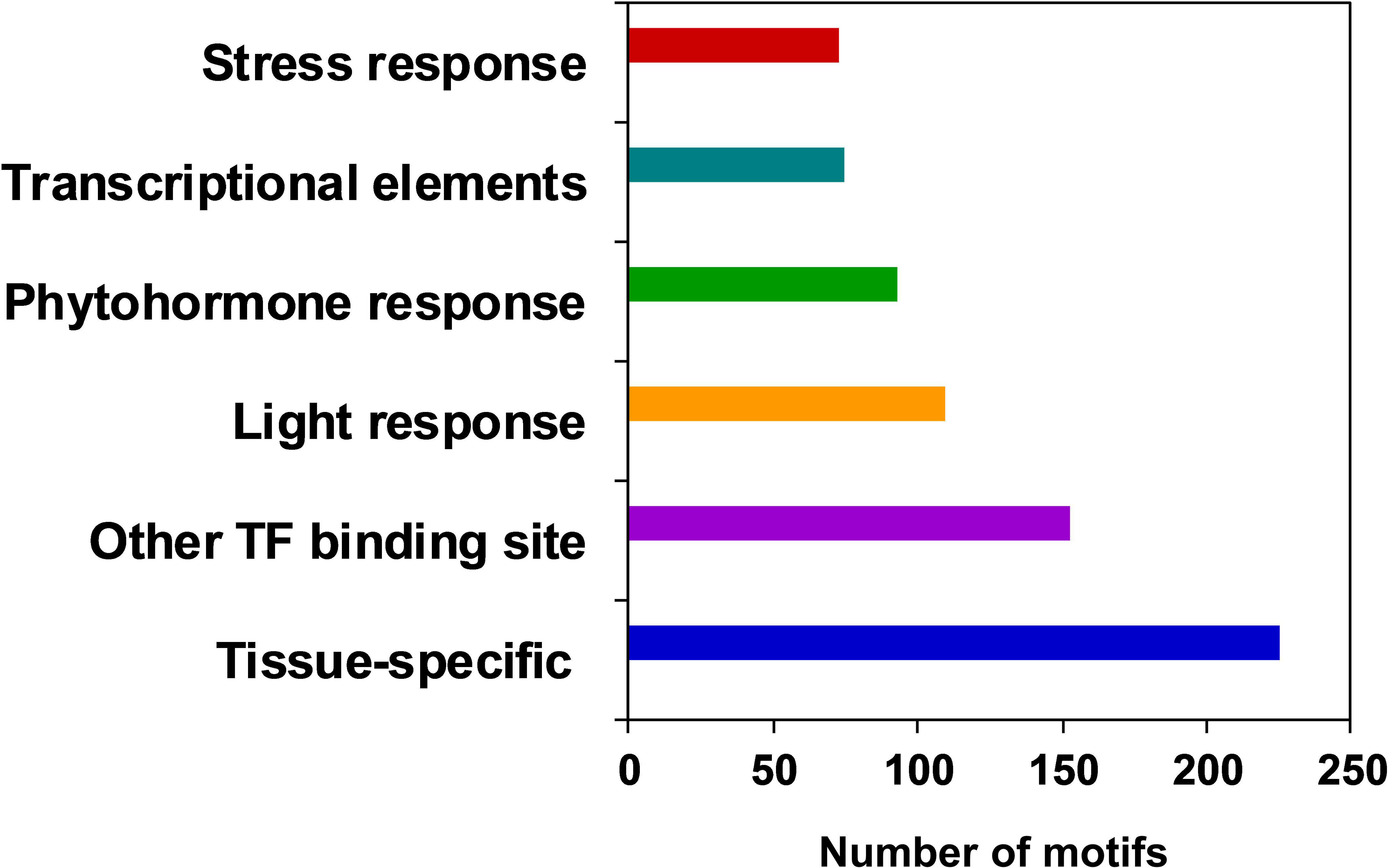
Functional classification of *cis*-elements present in the *GhNAC4* promoter. The motifs in the GhNAC4 promoter were grouped into five functional categories. There are a relatively high proportion of tissue-specific motifs and transcription factors binding sites.

**Table 2.**
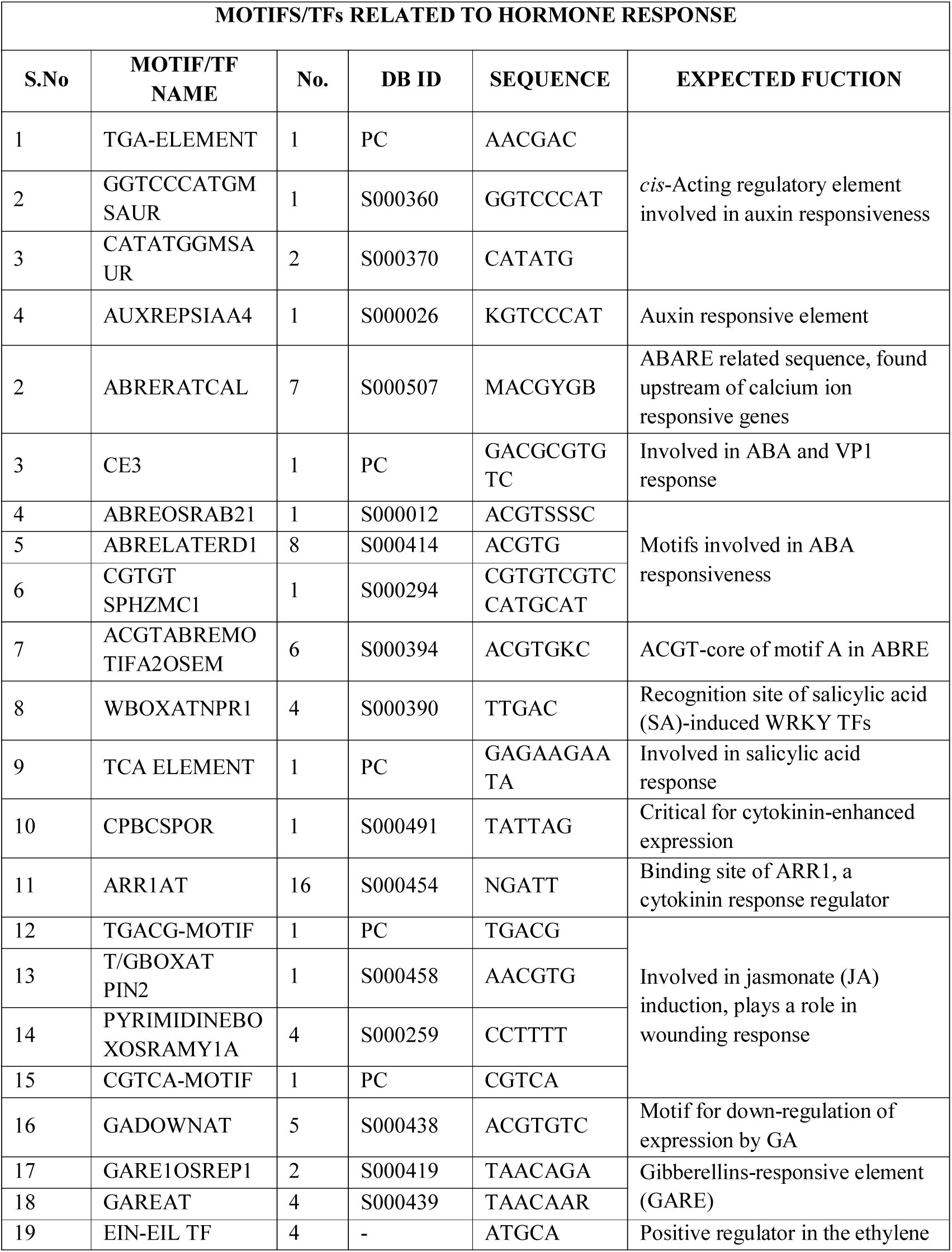
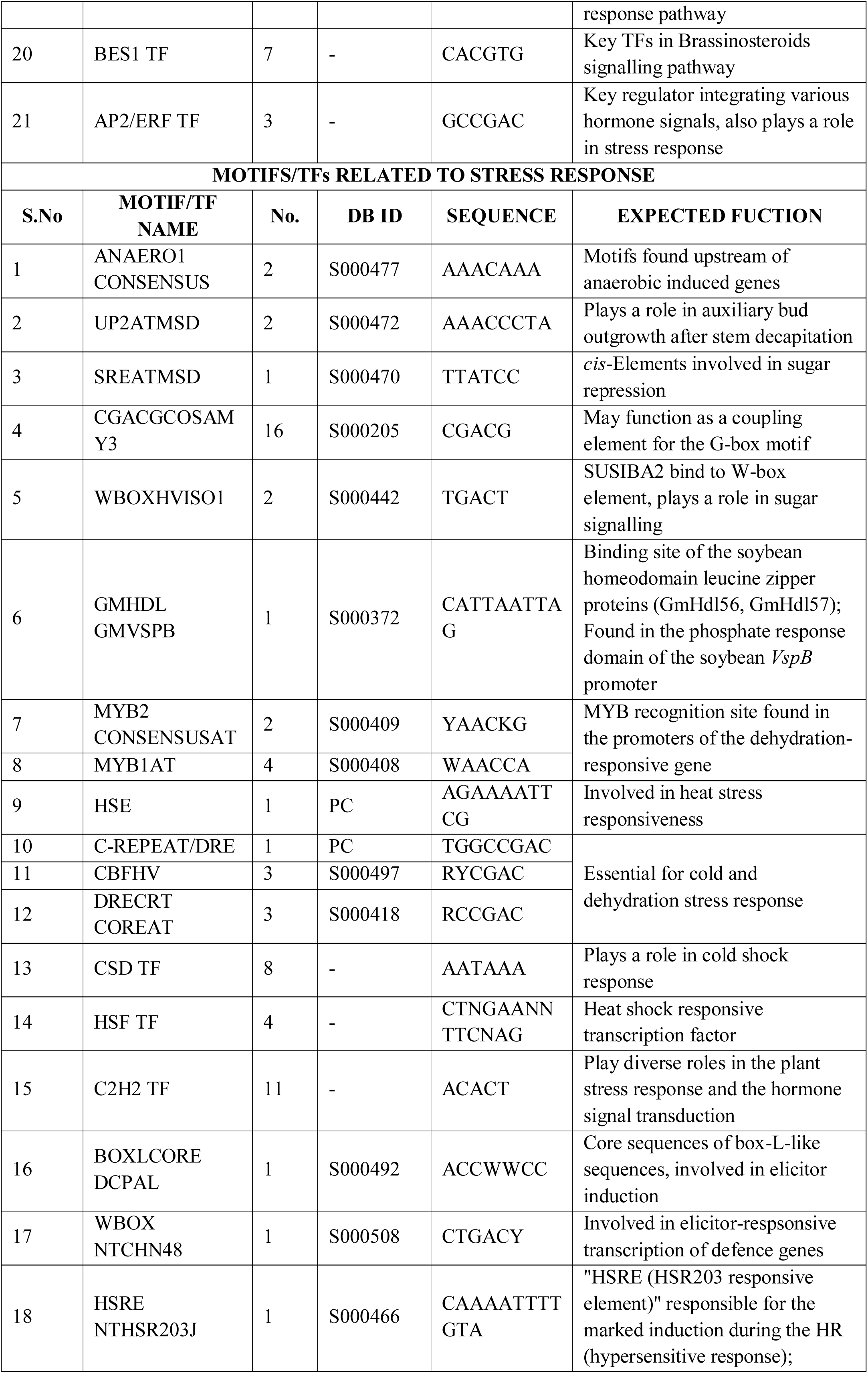
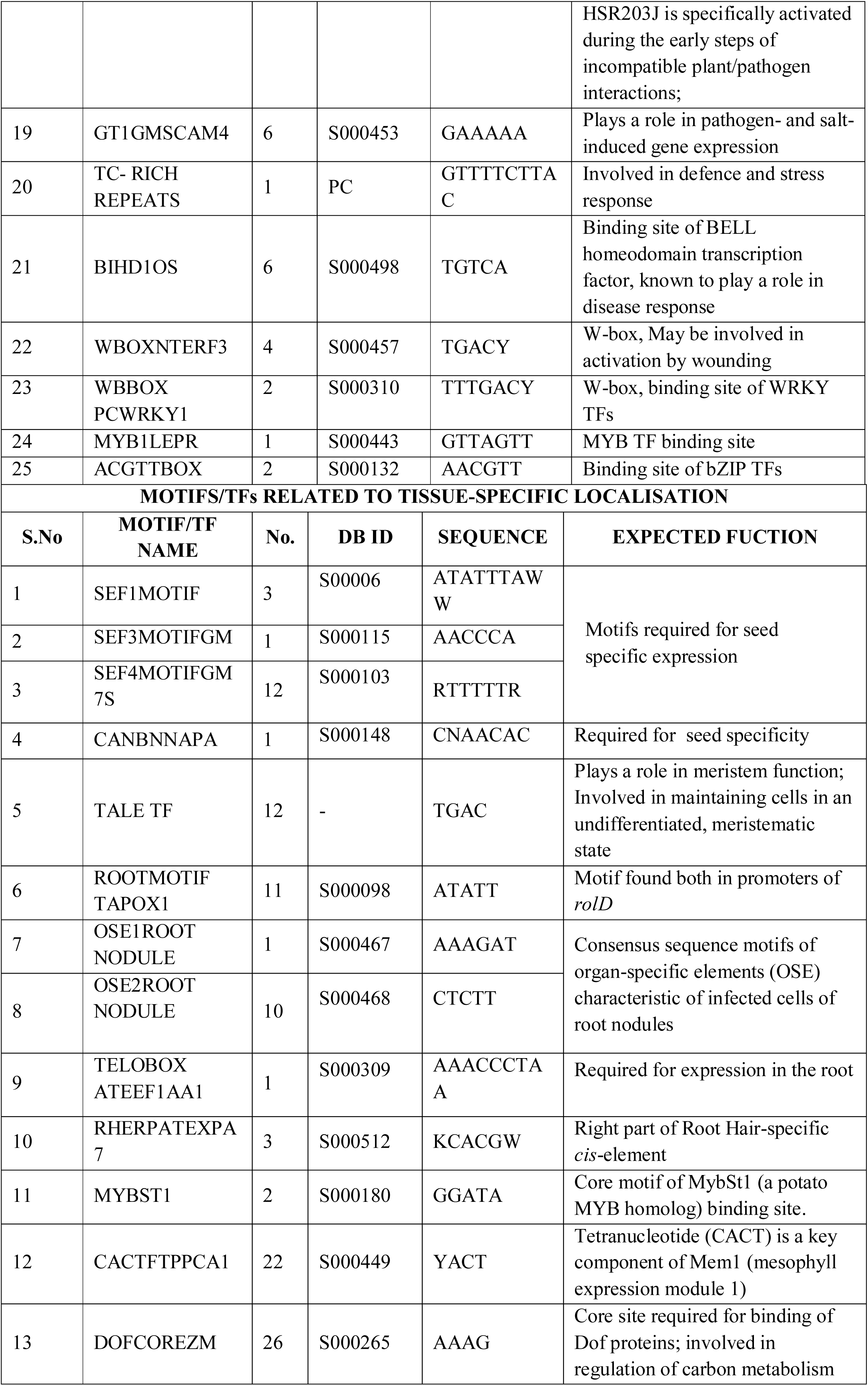
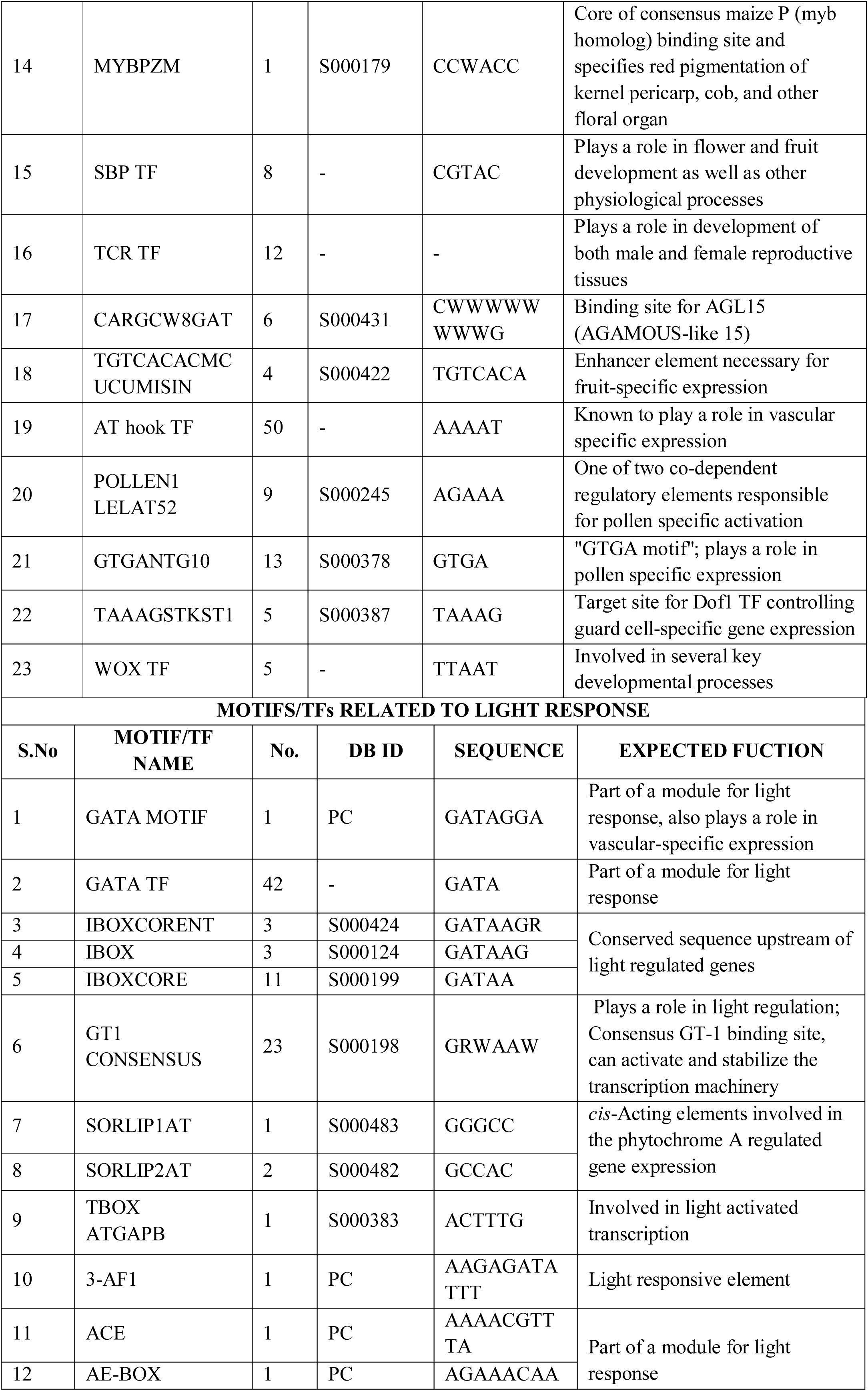
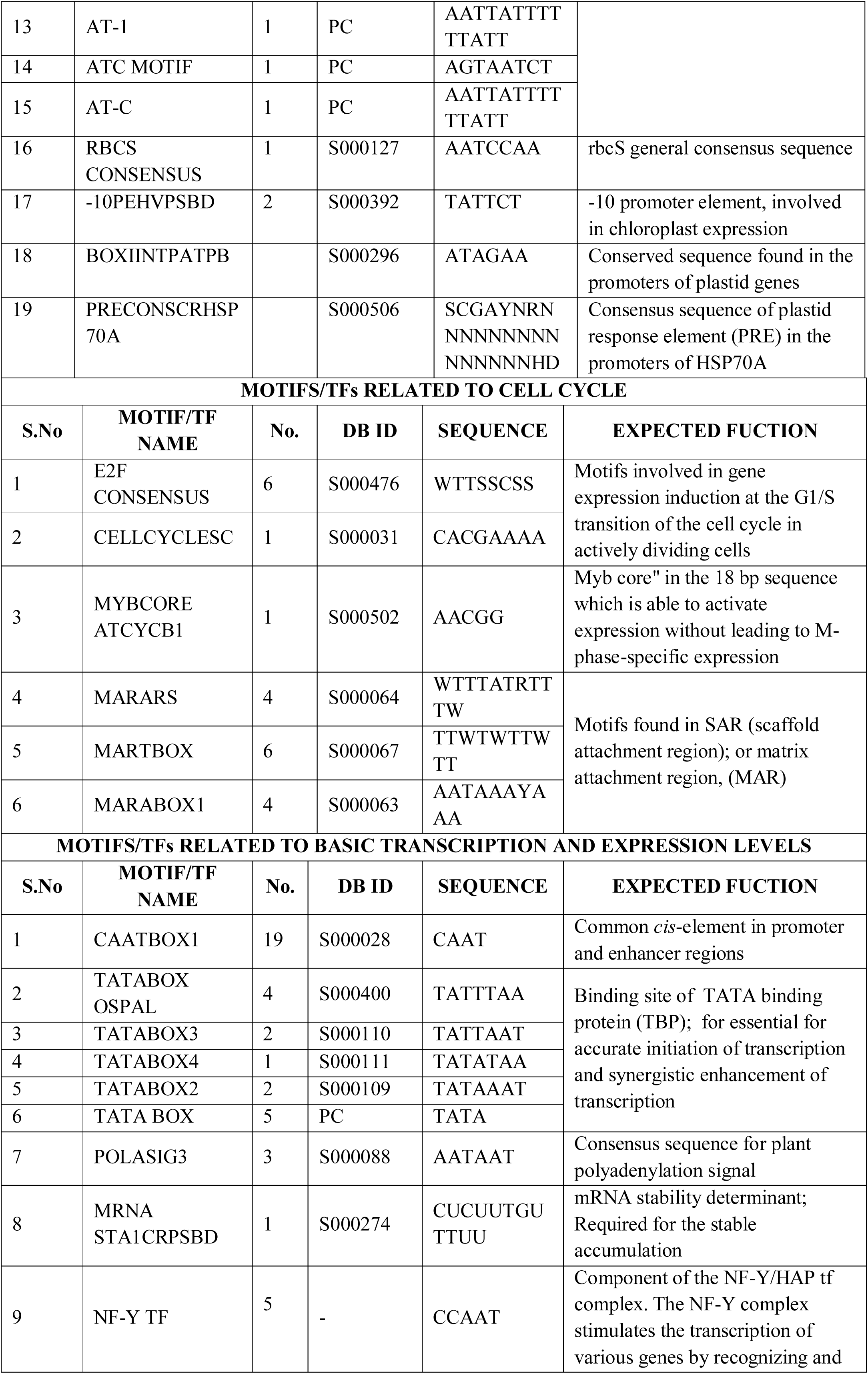
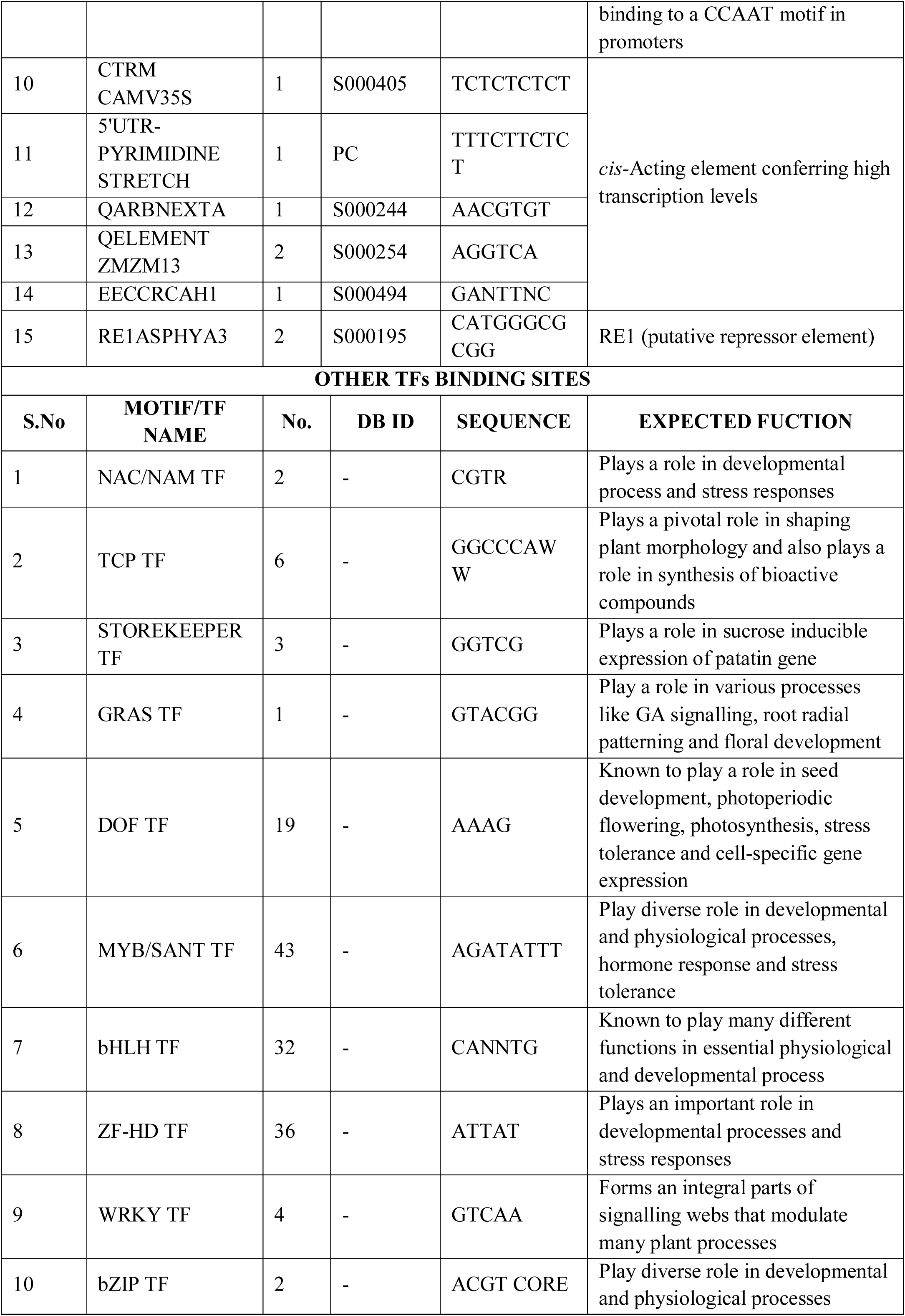
Putative motifs in the *GhNAC4* promoter identified by *in silico* analysis. DB ID represents the PLACE database accession number. PC represents the motifs identified by PlantCare database

### Hormone-related *cis*-acting elements predicted on *GhNAC4* promoter sequence

Several motifs responding to phytohormones were revealed in the *GhNAC4* promoter. Various ABA-responsive elements such as ABRELATERD1, ACGTABREMOTIFA2OSEM, and ABRERATCAL (found upstream of Ca^+2^ responsive genes) were observed in the promoter sequence. Auxin-responsive motifs such as one AUXREPSIAA4, one TGA element, one GGTCCCATGMSAUR, and two CATATGGMSAUR motifs, were predicted. Salicylic acid responsive motifs like TCA element and WBOXATNPR1 (recognition site of SA-induced WRKY TFs) were also identified. TGACG and CGTCA motifs required for jasmonic acid induction were found in the sequence. Gibberellin responsive elements such as GARE1OSREP1, GAREAT, and GADOWNAT were found on the putative promoter sequence.

ARR1AT TF, a cytokinin response regulator, and CPBCSPOR motif critical for cytokinin enhanced expression were observed. The *cis*-regulatory elements known to play a role in ethylene induction such as ERELEE4 and binding sites of EIN/EIL TF, a positive regulator in the ethylene response pathway were also identified on the promoter sequence. The binding sites for BES1 TF, a key TF in the Brassinosteroids signalling pathway were also observed in the putative promoter region. Furthermore, several recognition sites of AP2/ERF TFs, which are key regulators in integrating various hormone signals, were also found in the sequence and play roles in stress responses.

### Stress regulating *cis*-acting elements on the promoter sequence

A scan of the motifs showed that some potential MYB recognition sites like MYB1AT and MYB2CONSENSUSAT that are associated with stress regulation are observed in the promoters of the dehydration-responsive genes. *cis*-Regulatory elements such as DRECRTCORE, HSE, CBFHV and a binding site for CSD and HSF TFs, known to regulate responses to cold shock and heat stress were predicted. The binding site for bZIP (ACGTTBOX), WRKY (WBBOXPCWRKY1) and MYB (MYB1LEPR) TFs that were known to play important roles in stress response were found on the putative promoter region of *GhNAC4*. *GhNAC4* promoter sequence also contained a *cis*-acting element for nutrient deficiency response like the GMHDLGMVSPB, which is a binding site of homeodomain-leucine zipper protein, which is found in the phosphate response domain.

Various motifs and target sites of TFs, vital for biotic stress response were also predicted in the promoter sequence of *GhNAC4*. Many fungal elicitor response elements like BOXLCOREDCPAL, TC-RICH REPEATS, and GT1GMSCAM4 were also observed. Numerous binding sites of BELL homeodomain TF, BIHD1OS that is known to play a vital role in disease response were also observed in the sequence. Hypersensitive response element like HSRENTHSR203J was revealed. Motifs involved in wounding response such as T/GBOXATPIN2 and WBOXNTERF3 were also found. ANAERO1CONSENSUS, a motif found upstream of anaerobic induced genes was identified.

### Sugar and light-responsive motifs were predicted on *GhNAC4* promoter sequence

The *cis*-acting elements involved in sugar repression like PYRIMIDINEBOXOSRAMY1A, UP2ATMSD and WBOXHVISO1 (SUSIBA2 binds to W-box element) TFs were identified; they are also known to play a role in auxiliary bud outgrowth after stem decapitation.

Several motifs essential for light-regulated transcriptional activation like SORLIP2AT, IBOXCORE, GT1CONSENSUS (that activate and stabilise the transcription machinery), GATA (that also plays a role in the vascular-specific expression) and CGACGOSAMY3 (G-Box/BOXII) were identified in the putative promoter sequence suggesting that *GhNAC4* might be highly regulated by light. Motifs important for plastid-specific expression such as BOXIINTPATPB, -10PEHVPSBD, and PRECONSCRHSP70A were also present in the *GhNAC4* promoter.

### *cis*-Acting elements controlling tissue-specific expression

The specific expression is crucial for genes functioning at particular stages and in a particular tissue(s). A plethora of motifs essential for tissue-specific expression was predicted in the *GhNAC4* promoter sequence. Motifs important for embryo maturation and seed development such as SEF4MOTIFGM7S, SEF1MOTIF, and CANBNNAPA were found. Binding sites for TALE TFs, which are known to a play role in meristem function and involved in maintaining cells in an undifferentiated state, were also observed. Motifs involved in root specific expression like OSE2ROOTNODULE, ROOTMOTIFTAPOX and recognition sites for MYBST1 TF were identified.

Several copies of CACTFTPPCA1, which is an important motif for mesophyll cell expression, were observed in the sequence. Numerous binding sites of DOFCOREZM involved in the regulation of carbon metabolism and shoot and leaf-specific expression were identified in the promoter sequence. Binding sites of WOX TF, known to be involved in several key developmental processes was also observed. TAAAGSTKST1, a motif known to play a role in controlling guard cell-specific gene expression was also observed. Numerous binding sites for TFs significant for vasculature-specific expression like DOF, GATA and AT hook TFs were observed in the putative promoter sequence. Many target sites for TCR, SBP and MYBPZMTFs and motifs like CARGCW8GAT and TGTCACACMCUCUMISIN significant for flower and fruit development were present. Motifs for pollen specific expression such as POLLEN1LELAT52 and GTGANTG10 were also identified.

### Cell-cycle specific and basal TF binding elements predicted on the promoter sequence

*cis*-Acting elements involved in gene expression induction in actively dividing cells such as MYBCOREATCYCB1, E2FCONSENSUS and CELLCYCLESC were predicted in the sequence. Motifs such as MARARS, MARABOX1, and MARTBOX are known to play a role in scaffold attachment region were also found.

Several potential binding sites for transcription factors, such as NAC, MYB/SANT, ZF-HD, bHLH, Storekeeper, bZIP, GRAS, and TCP were also identified.

Basal motifs that play a critical role in the core transcription initiation were identified in the promoter of *GhNAC4*. Several TATA box motifs, essential for accurate initiation of transcription and synergistic enhancement of transcription were found. CAAT-box motifs that are universal enhancer elements in promoters were also predicted. In addition to basal regulatory elements, many transcriptional regulatory elements were found. Several enhancer elements viz., CTRMCAMV35S, QARBNEXTA, QELEMENTZMZM13, and 5’ UTR-PYRIMIDINE STRETCH and binding sites of NF-Y TF were identified. Interestingly, a repressor element like RE1ASPHYA3 was also found in the promoter. These elements suggest that *GhNAC4* promoter is functional *in situ*.

The presence of these motifs provide hints on the regulation of *GhNAC4* gene expression in cotton as complex and it may play an important role during plant growth and development and is responsive to phytohormones, environmental stresses, and light.

### Generation and analysis of tobacco transgenics of PRO*_GhNAC4_*:GUS

To evaluate the promoter activity of *GhNAC4* gene, the 1612 bp fragment was amplified by using appropriate PCR primers and the resultant fragment was cloned into pTZ57R/T vector for confirmation through sequencing. The promoter sequence was then cloned into a plant binary vector pCAMBIA1381Z for fusing it with the *uidA* (GUS) reporter gene in a transcriptional fusion. Transgenic tobacco plants carrying the PRO*_GhNAC4_*:GUS were generated using the standard Agrobacterium-mediated leaf disc transformation method. A total of 12 hygromycin resistant T_0_ plants were generated and were confirmed by genomic PCR. Out of these, 11 plants exhibited the expected band sizes of 1612 bp and 1026 bp corresponding to the size of *GhNAC4* promoter sequence and the *HptII* gene, respectively. To eliminate the effect of gene copy number on GUS activity, only single copy T_1_ progenies, P7, P9, and P17 were used for further generation of T_2_ seeds. The homozygosity of T_2_ lines was confirmed by uniform 100% germination on a medium supplemented with the inhibitory concentration of hygromycin. The T_2_ progenies of three lines, P7.1, P9.5, and P17.3 were used for tissue-specific localisation and MUG assay.

### *GhNAC4* localizes to various tissues during growth and development

As several motifs pertaining to tissue localisation were observed in the promoter sequence, we used GUS histochemical assay to precisely define the spatio-temporal expression patterns of *GhNAC4* promoter throughout plant development in T_2_ generation transgenic tobacco plants. Fig. 5 shows the localisation of PRO*_GhNAC4_*:GUS fusion in vegetative and reproductive tissues in transgenic tobacco plants.

**Fig. 5.**
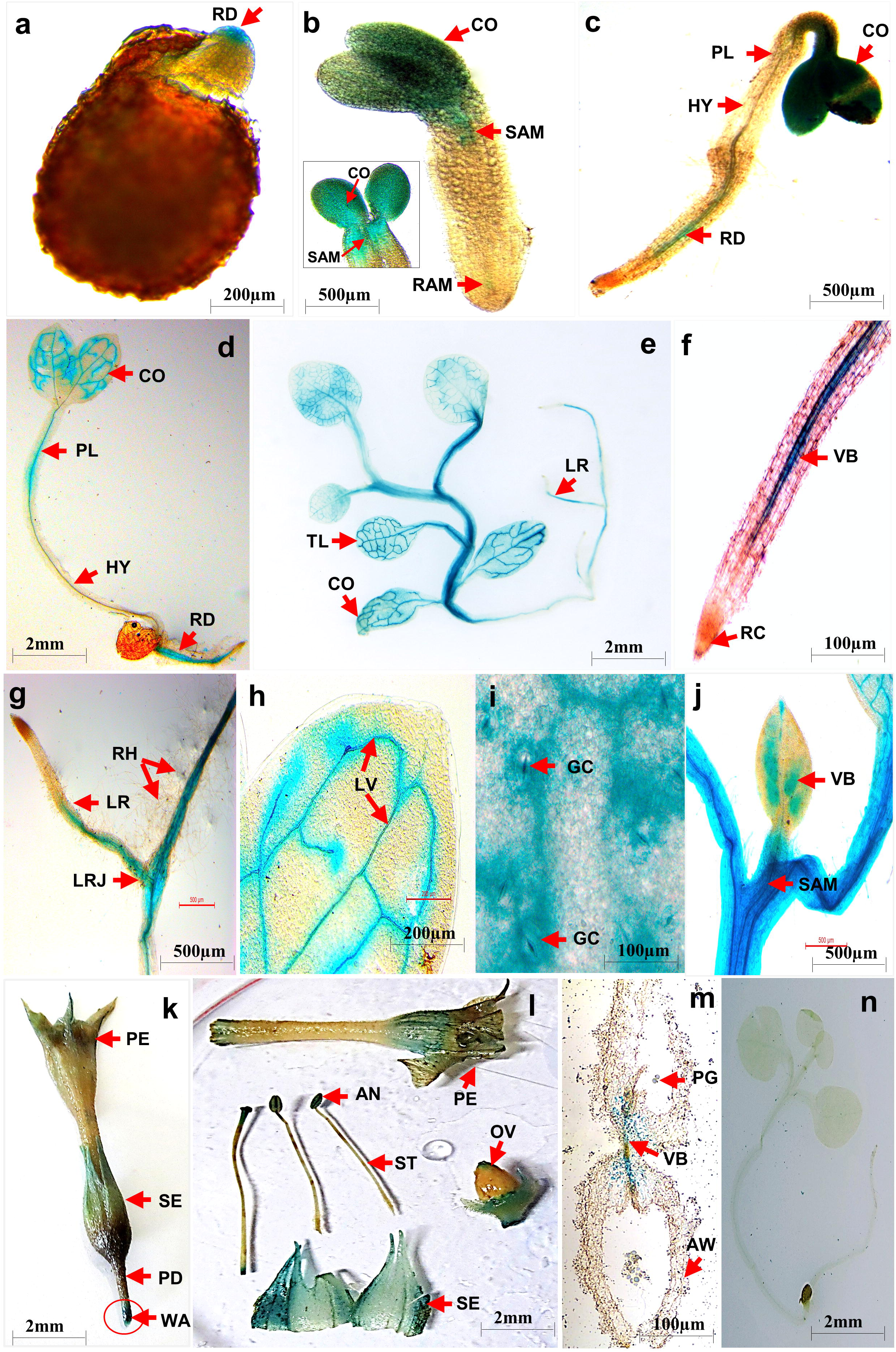
Histochemical localisation of GUS activity in tobacco transgenic plants containing PRO*_GhNAC4_*:GUS construct. **a**-**e** Seedlings grown on MS media with hygromycin at - **a** Day1; **b** Day 3; **c** Day 7; **d** Day 15; **e** Day 30. **f**-**j** Various tissues of a 30-day old transgenic tobacco plant - **f** Main root; **g** Lateral root; **h** True leaf; **i** Guard cells; **j** Developing leaf. **k**-**m** Floral structures – **k** Mature flower; **l** Dissected flower showing various tissues; **m** Cross-section of an anther. **n** 15 days old tobacco seedling carrying empty pCAMBIA 1381Z vector. AW, anther wall; CO, cotyledon; GC, guard cell; HY, hypocotyl; LR, lateral root; LRJ, lateral root junction; LV, lateral vein; OV, ovary; PD, pedicel; PE, petal; PG, pollen grains; PL, plumule; RAM, root apical meristem; RD, radicle; RH, root hairs; SAM, shoot apical meristem; SE, sepal; TL, true leaf; VB, vascular bundle; WA, wounded area. All arrows show strong GUS activity or no activity. Bars of each panel are as shown

In the early stages of tobacco growth (1 d old seedlings), GUS activity was first observed in the emerging radical (Fig. 5a), which was later detected in emerging cotyledons, root tip and shoot apex of 3 d old seedlings. However, the GUS activity was relatively weaker in the hypocotyl tissue (Fig. 5b). In 7, and 15 d old transformed tobacco seedling, GUS expression was detected in the leaf veins, petioles, stem and root (Fig. 5c-d). Similar GUS activities were maintained in the one-month-old plants (Fig. 5e).

The main and lateral roots showed GUS expression, which was absent in the root cap region and the root hairs (Fig. 5f-g). Intense GUS activity was also observed in the mid rib and the lateral veins, but the leaf lamina showed little staining (Fig. 5 h). We also observed the GUS activity in the guard cells (Fig. 5i) and the developing mid rib regions of a young leaf (Fig. 5j). Fig. 5k-m shows the staining of the floral structures, which revealed that the GUS activity was present in sepals, anthers, pollen grains, and the stigma and to a lesser extent in the petal edges. However, it was absent in the ovary and pedicel. No staining was detected in the seedlings harbouring a promoter-less *GUS* gene regardless of the developmental stage (Fig. 5n). This strongly indicates that *GhNAC4* expression is under developmental and temporal control, and its promoter modulates a precise transcriptional regulation throughout the plant growth and development processes, which are consistent with the *cis*-regulatory elements predicted on it.

### *GhNAC4* showed strong expression in vascular bundles and is wound-inducible

To obtain a better understanding of tissue specificity of GUS activity, thin cross-sections of various tissues using a razor blade were made. Strong GUS staining was detected only in the vascular bundles, especially in the phloem of leaf veins, petiole, stem, and root (Fig. 6a-e). Both the abaxial and adaxial phloem showed intense GUS staining. Other cell types, like the pith, cortex, and epidermis remained unstained. However, upon wounding, cortex, and epidermis also showed intense GUS expression (Figs. 5k and 6f).

**Fig. 6.**
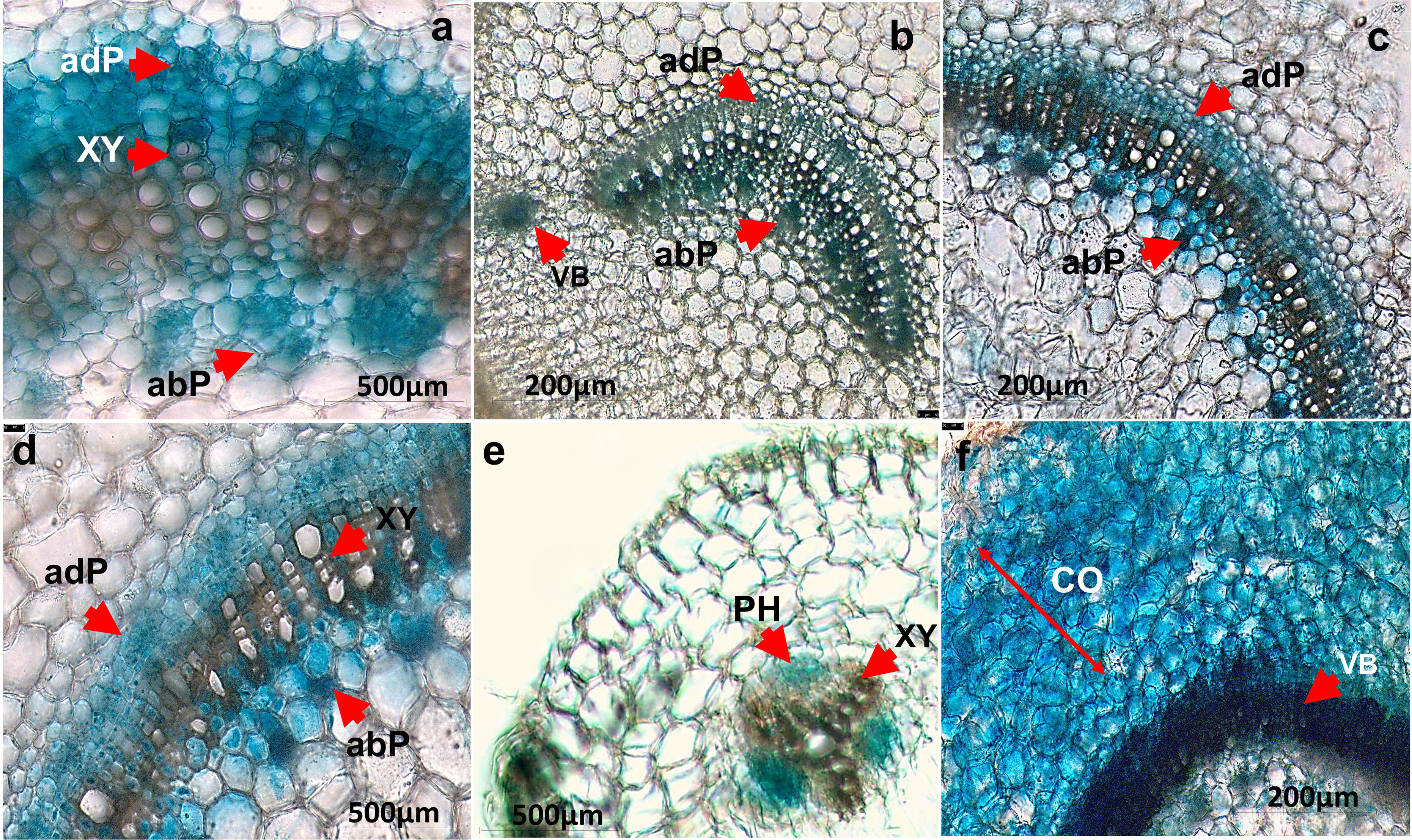
GUS activity in free hand cross-sections tobacco transgenic plants containing PRO*_GhNAC4_*:GUS construct. **a** Leaf; **b** Petiole; **c** Stem **d** Magnified view of the stem; **e** Root; **f** Wounded stem. abP, abaxial phloem; adP, adaxial phloem; CO, cortex; P, phloem; VB, vascular bundle; XY, xylem. All arrows show strong GUS activity or no activity. Bars of each panel are as shown

### *GhNAC4* promoter is induced by various phytohormones in transgenic tobacco seedlings

To explore the possible regulation of *GhNAC4* promoter by phytohormones, the activity of GUS was examined by fluorometric MUG assay, and treating the T_2_ transgenic tobacco seedlings expressing the PRO*_GhNAC4_*:GUS fusion with various hormones. The corresponding results have been depicted in Fig. 7. The specific activity of GUS enzyme without any hormone treatments in GhNAC4 seedlings was measured to be 2.42±0.42pmol µg^−1^ min^−1^. This is in agreement with the histochemical staining results, where GUS staining was observed even in untreated seedlings. When 2-week old tobacco seedlings were treated with different hormones for 24 h, the highest level of induction was seen upon BAP treatment (∼3.1 fold) and followed by ABA (∼2.7 fold). GUS activities were also moderately enhanced by IAA (∼2.4 fold) and MeJA (∼2.2 fold). The promoter weakly responded also to GA3 (∼2.1 fold) and Ethylene (∼1.8 fold), but not to SA as compared with the untreated control samples. This suggests that *GhNAC4* promoter is regulated largely by multiple phytohormones. This data is consistent with the pattern and presence of *cis*-acting elements predicted in the promoter sequence.

**Fig. 7.**
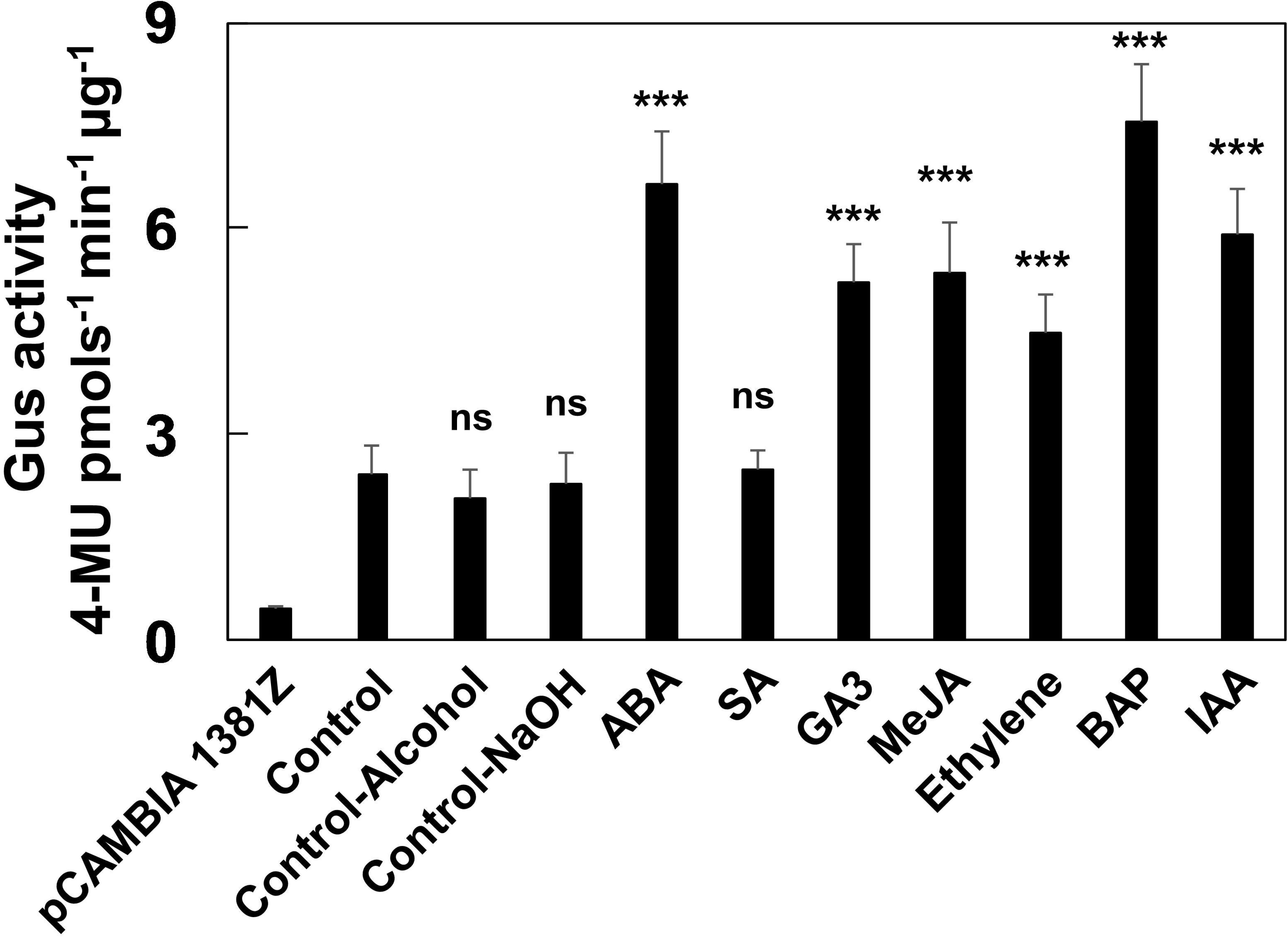
Effect of various phytohormones on the GUS activity of PRO*_GhNAC4_*:GUS tobacco transgenics. Fluorometric analysis of the *GUS* enzyme in tobacco seedlings after ABA, MeJA, SA, 6-BAP, GA_3_, IAA, or Ethephon treatment. Two weeks-old PRO*_GhNAC4_*:GUS tobacco transgenic seedlings incubated for 24 h under the treatment, were used for analysis. pCAMBIA 1381Z empty vector harbouring tobacco seedlings were used as a negative control. The data are shown as the means ± SE (n=3). A statistical analysis with one-way ANOVA indicates significant differences (*** P<0.001, ns - not significant)

### *GhNAC4* promoter is responsive to various environmental stress treatments

Since we have observed that *GhNAC4* gene expression was regulated by various stresses, we studied the promoter activity in transgenic tobacco plants harbouring the PRO*_GhNAC4_*:GUS fusion by treating them with various environmental stresses for varying time points. This study was undertaken after taking a cue from the presence of several stress-responsive motifs in the promoter sequence, and the GUS activity was measured by the fluorometric MUG assay. An approximately three-fold increase in GUS activity was observed when the seedlings were subjected to salt, mannitol, PEG, MV, and air-drying as compared to untreated seedlings. A combination of cold and dark treatment also activated the promoter (∼2.4 fold) as compared to dark alone (∼1.7 fold). Flooding stress caused by submerging the seedlings in water moderately induced the promoter (∼1.9 fold). High temperatures and wounding activated the promoter weakly (∼1.7 fold) as shown in Fig. 8.

**Fig. 8.**
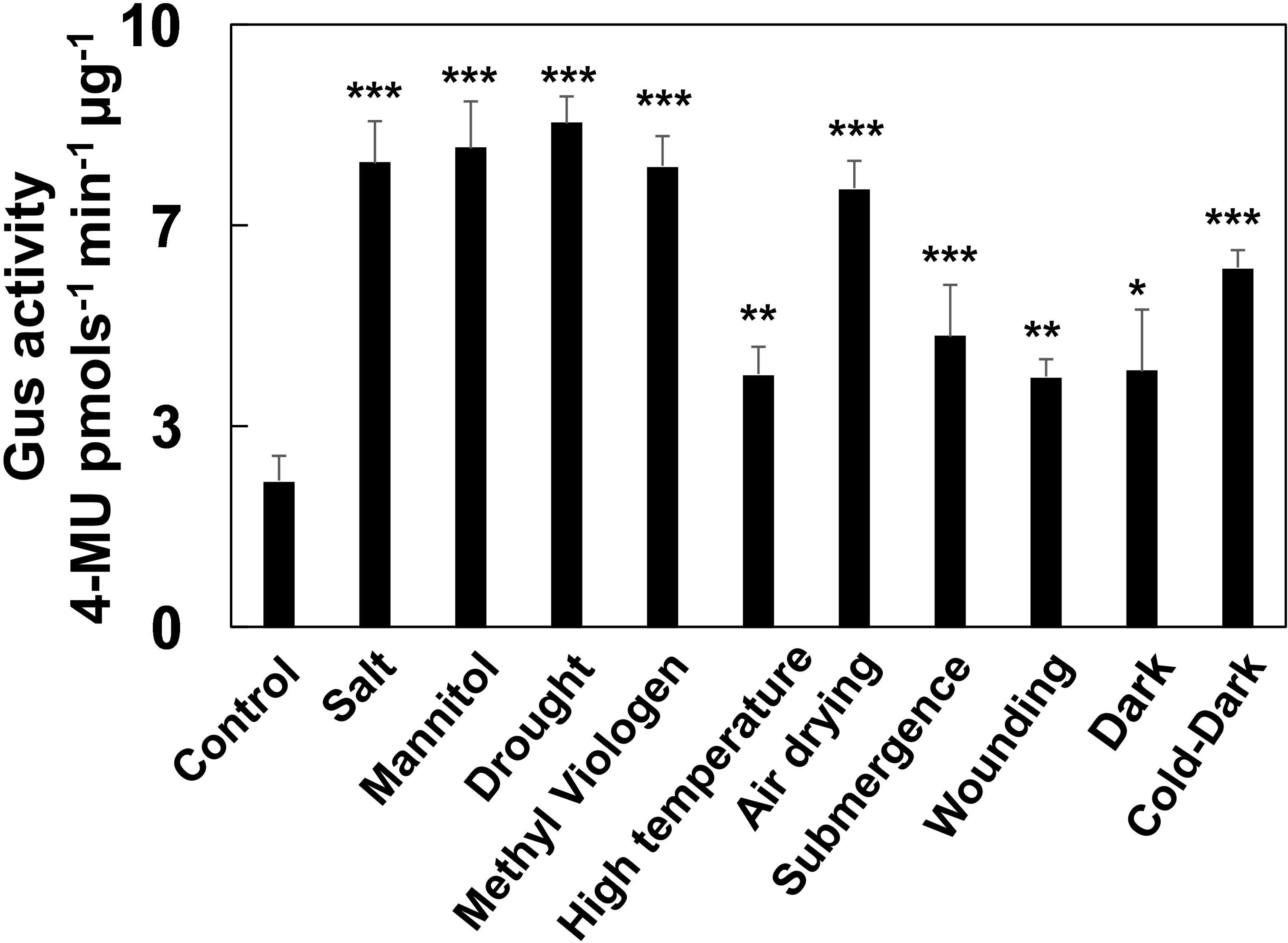
Effect of various stresses on the GUS activity of PRO*_GhNAC4_*:GUS tobacco transgenics.Fluorometric analysis of the *GUS* enzyme in tobacco seedlings after NaCl, mannitol, PEG, methyl viologen, high temperature, air drying, submergence, wounding, dark or combination of dark and cold treatment. Two weeks-old PRO*_GhNAC4_*:GUS tobacco transgenic seedlings, were used for analysis. The data are shown as the means ± SE (n=3). A statistical analysis with one-way ANOVA indicates significant differences (* P<0.05, ** P<0.01, *** P<0.001)

There are some differences in the expression and induction levels among experimental systems such as cotton seedlings and tobacco transgenics, which suggest the presence of other factors in the respective experiments in terms of treatments.

## Discussion

Transcriptional regulation is essential for mediating the responses of an organism to environmental cues (López-Maury et al. 2008). The promoter of a gene is critical in determining its spatial and temporal expression in the plant and during its development (Singh 1998). The promoter sequence specifies the recruitment of TFs, which regulate its pattern of gene expression. The short *cis*-acting elements that specify the binding for the TFs are the vital functional components of promoter function. Therefore, investigations into the *cis*-acting elements and their interactions would aid in the understanding of the mechanism of transcriptional regulation (Hudson and Quail 2003).

To identify the putative *cis*-regulatory elements, a motif search was carried out for the promoter sequence of *GhNAC4*. We found that *GhNAC4* promoter region contained several *cis*-acting elements linked to various stress responses, phytohormone induction, tissue-specific localisation, sugar response, light response, and transcriptional activation.

To precisely evaluate the function of the *GhNAC4* promoter, we generated transgenic tobacco plants expressing the PRO*_GhNAC4_*:GUS construct and analysed the spatial, temporal, and developmental expression of GUS. We also studied the *GhNAC4* expression and promoter induction by GUS enzyme activity in treatments with the various external phytohormones and environmental stressors.

### *GhNAC4* promoter exhibits hormone responsiveness

Understanding how hormones and genes interact in a changing environment to synchronise plant growth and development is an important breakthrough in plant developmental biology (Vanstraelen and Benková 2012). In a stressed condition, phytohormones might exhibit either synergistic or antagonistic interactions to modulate plant growth (Robert-Seilanian et al. 2011). Several phytohormones such as GA3, BAP, ABA, and MeJA induced the expression of *GhNAC4* by 7-11 folds. The treatment with other hormones such as ethylene, SA and IAA led to a moderate increase in expression levels (4-5 fold). *GhNAC4* promoter region carries several key motifs that could be linked to the observed hormone responses.

ABA is an important stress hormone that plays crucial roles in plant adaptation to different stress conditions like osmotic imbalance, salinity, and drought (Vishwakarma et al. 2017). It regulates the expression of many genes that have functions in abiotic stress tolerance (Fujita et al. 2011). *GhNAC4* shows several folds induction by ABA (Figs. 1 and 7) and its promoter carried several motifs known for ABA responsiveness such as six ACGTABREMOTIFA2OSEM, eight ABRELATERD1 and one ABREOSRAB21 motifs. ACGTABREMOTIFA2OSEM is an ABA-responsive motif found in the promoter of the rice *OsEm* gene, which is regulated by the seed-specific VP1 TF (Hattori et al. 2002). The ABRELATERD1 motif is related to the ABRE motif, which has been found upstream of Early Response to Dehydration1 (*ERD1*) gene and is responsive to ABA showing significant up-regulation under water stress (Simpson et al. 2003). A single copy of ABRE is insufficient for ABA-responsive regulation of gene expression. Either multiple copies of ABRE motif or ABRE along with a coupling element forming the active ABA Response Complex (ABRC) is essential for ABA-induced gene expression (Skriver et al. 1991). *GhNAC4* promoter has been predicted to have one *CE3* coupling element. The minimal ABRC of the *HVA1* promoter from barley exhibited one coupling element and along with one ABRE motif, which is enough to confer high levels of ABA induction when linked to a minimal promoter (Shen et al. 1996).

Regulation of gene expression by cellular calcium plays a crucial role in plant responses to environmental stresses (Reddy et al. 2011). The stress stimuli trigger a burst of cytosolic calcium ions, which confers changes in gene expression, thereby allowing plants to adapt to the stress conditions (Knight 2000). The ABRERATCAL motif is a sequence related to ABRE that is found upstream of calcium ion responsive genes. Quite a few genes upregulated by calcium ion exhibiting ABRERATCAL motifs were early stress-induced genes (Kaplan et al. 2006). The *GhNAC4* promoter region was predicted to have seven ABRERATCAL motifs suggesting that it may also be responsive to calcium and ABA-mediated stress adaptation.

Gibberellins (GA) are essential for many plant developmental processes such as seed germination, plant growth and flower development (Richards et al. 2001). GA and ABA are known to mediate several plant developmental processes antagonistically and the recent data suggests the involvement of GA in plant adaptation to stress (Colebrook et al. 2014). *GhNAC4* is highly upregulated by GA3, and its promoter has four copies GAREAT and two copies of GARE1OSREP1 motifs, which are gibberellin responsive elements (Skriver et al. 1991). An R2R3-type MYB TF called the GAMYB directly interacts with the GARE motif in the promoter of barley amylase gene and seemed to be essential for the GA induced gene expression (Gubler et al. 1995). Subsequent evidence indicated that GAMYB is also a target of the antagonistic effects of ABA signalling (Gomez-Cadenas et al. 2001; Zentella et al. 2002). Multiple copies of GARE alone or in association with other motifs such as PYRIMIDINE box or TATCCAC box (forming the Gibberellins Response Complex) are essential for GA induced gene expression at higher levels (Lanahan et al. 1992; Rogers et al. 1994). *GhNAC4* promoter region also exhibits four copies of PYRIMIDINEBOXOSRAMY1A motif. A DOF TF (OsDOF3) binds to the pyrimidine box in the promoter region of rice *RAMY1a* gene that is one of the most prominent GA-responsive genes (Washio 2001). GAMYB and OsDOF3 are shown to interact with each other and synergistically modulate GA induced gene expression (Washio 2003). GA is the main target for stress-induced growth modulation and recent evidence also indicates the involvement of GA in either promotion or suppression of growth depending on the type of stress (Colebrook et al. 2014). GADOWNAT motif is a common sequence found in the genes down-regulated by GA. Interestingly, this motif is identical to ABRE (Ogawa et al. 2003). Five copies of this motif are present in the *GhNAC4* promoter suggesting coordinated regulation of *GhNAC4* by GA and ABA.

Ethylene is another hormone essential for plant growth and development playing a crucial role in responses to environmental stress such as pathogen attack, wounding, flooding, high temperatures, and drought (Abiri et al. 2017). *GhNAC4* promoter showed the presence of four binding sites for ethylene insensitive (EIN) TF, and the ethylene treatment upregulated the transcript levels of *GhNAC4*. AtNAC2 TF up-regulation under salinity treatment was induced in ethylene-overproducing mutant *eto1-1* but was repressed in ethylene-insensitive mutants *etr1-1* and *ein2-1* (He et al. 2005) suggesting that ethylene plays a positive role in the salt response of AtNAC2 TF. Most of the ethylene responses appear to be mediated by EIN3 TF along with ethylene insensitive-like (EIL) protein (Solano et al. 1998). EIN3 TF was shown to enhance salt tolerance in Arabidopsis (Peng et al. 2014) and also act synergistically with SOS2 to modulate salt stress response (Quan et al. 2017). Furthermore, EIN3/EIL1 act as a node integrating ethylene and JA signalling for regulating plant growth and stress responses (Zhu et al. 2011).

Cytokinin regulates many important aspects of plant growth and development such as the development of vasculature, photomorphogenesis and stress responses (Werner and Schmülling 2009). We have observed high up-regulation of *GhNAC4* transcripts in cytokinin (BAP) treatment, and the promoter region of this gene evidenced one CPBCSPOR motif and 16 binding sites for Authentic Response Regulators1 (ARR1). CPBCSPOR motif was found upstream of cucumber NADPH-protochlorophyllide reductase (*POR*) gene and is essential for cytokinin dependant transcriptional activation, which is involved in interactions with cytokinin related elements (Fusada et al. 2005). ARR1 is an important signalling component acting at the head of transcriptional cascade to regulate cytokinin response and is known to be involved in cytokinin-mediated differentiation of protoxylem (Yokoyama et al. 2007). They are also known to regulate plant responses to abiotic stresses (Nguyen et al. 2016; Jeon and Kim 2013).

Auxin is a phytohormone that plays a crucial role in growth and development. It regulates the development of primary and lateral roots and has a most profound role in vascular differentiation and venation pattern formation (Zhao 2010). Auxin (IAA) was shown to induce the expression of *GhNAC4* and the promoter region shows an AUXREPSIAA4, one TGA element, one GGTCCCATGMSAUR, and two CATATGGMSAUR motifs. AUXREPSIAA4 is an auxin-responsive element (AUXRE) found upstream of pea *IAA4* gene, which is an early auxin-inducible gene (Ballas et al. 1993). GGTCCCATGMSAUR and CATATGGMSAUR are AUXREs found in the promoter region of soybean *SAUR* (Small Auxin-Up RNA) gene (Li et al. 1994; Xu et al. 1997). Each of these promoters is rapidly (within a few minutes) and specifically induced by auxin. Auxin Response Factors (ARFs) bind to AUXRE and mediate hormone responses. ARF5 is required for the formation of vascular strands at all stages and critically required for embryonic root formation (Hardtke and Berleth 1998). NAC1 place a key role in promoting auxin-mediated lateral root formation (Xie et al. 2000). Auxin role in stress response is emerging, and Bouzroud et al. (2018) suggested that ARFs are potential mediators of auxin action in response to environmental stresses. A plasma membrane-bound NAC TF NTM2 integrates auxin and salt signalling via the *IAA30* gene during seed germination in Arabidopsis (Park et al. 2011). Vascular bundle localisation of GUS in PRO*_GhNAC4_*:GUS transgenic plant could be attributed to the regulation of *GhNAC4* promoter by auxin and cytokinin.

Jasmonates are important regulators of plant defence responses to pathogen and insect attack, herbivory and wounding (Wasternack 2007). MeJA and wounding induced the expression of *GhNAC4* by several foldsand its promoter contained motifs such as T/GBOXATPIN2, TGACG, and CGTCA. JAMYC/AtMYC2 TF binds to the T/GBOXATPIN2 motif, found in the promoter of JA responsive and wound-inducible Protease Inhibitor II (*PIN2*) gene (Boter et al. 2004). JAMYC acts as a conserved master switch regulating the expression of several JA-regulated defence genes especially the genes involved in during wounding response (Lorenzo et al. 2004). ANAC019 and ANAC055 act downstream of AtMYC2 as transcriptional activators to regulate JA-signalled defence responses (Bu et al. 2008). A bZIP TF, TGA1 is a positive regulator of disease resistance and binds to the TGACG motif and its palindrome CGTCA motif which is known for JA responsiveness (Schindler et al. 1992; Shearer et al. 2012). TGACG motif is found in promoters of various JA responsive genes like *OsOPR1* that has been shown to play important roles in defence responses in rice (Sobajima et al. 2007).

Phytohormones act synergistically and antagonistically with each other to regulate plant growth and development in association with a changing environment by forming a complex cross-talks network (Robert-Seilaniantz et al. 2011). One of the TF families modulating these multiple regulatory responses is Apetala2/ Ethylene responsive factor (AP2/ERF) TFs (Licausi et al. 2013). Apart from ethylene, JA, ABA, auxin, and cytokinin also regulate many members of the AP2/ERF TF family (Phukan et al. 2017). AP2/ERF TFs, in turn, modulate the content of these phytohormones by regulating their biosynthesis pathways (Gu et al. 2017). The stimulated TFs would further regulate the downstream target genes resulting in changes in plant growth and development and environmental stress responses (Gu et al. 2017). They mediate downstream responses by binding to the GCC box (ethylene response element) and/or DREB element (Fujimoto et al. 2000). *GhNAC4* promoter has four copies of GCC box motif. Taken together, these data suggest that GhNAC4 may act downstream of AP2/ERF TF.

As *GhNAC4* expression is regulated by many phytohormones essential for stress response and plant development processes, and the promoter contains several motifs required for regulation by phytohormones. Our data suggest that GhNAC4 TF can act as a node modulating hormonal response in a changing environment to regulate plant development.

### Environmental stresses regulate GhNAC4

Plants being sessile, have developed capabilities to grow and propagate even under extreme environmental conditions such as high salt, severe drought, heavy metal stress, low or high temperatures (Zhu 2001). Plants have become specialized in rapidly sensing and responding to adverse environmental conditions by advancing a complex network of cellular processes for stress adaptation (Wang et al. 2003). Various genes induced during stress response not only play a role in stress tolerance, but also play a role in sensing and transcriptional regulation (Zhu 2016). *GhNAC4* was induced by drought stress (∼184 fold), salinity stress (∼43 fold), osmotic stress caused by mannitol (∼58 fold), oxidative stress induced by methyl viologen (∼14 fold), cold stress (∼19 fold), high temperature stress (∼6 fold), and wounding (∼5 fold) (Fig. 2). And the promoter was also induced by approximately 3 fold by salt, mannitol, PEG, MV, and air-drying stress treatments. Binding sites for various TF such as MYB, CSD, HSF, WRKY, BELL, and C2H2, known to be involved in environmental stress responses were observed in the *GhNAC4* promoter region (Gujjar et al. 2014; Singh et al. 2002).

Low-temperature stress and drought are two major limiting physiological factors that affect plant growth, productivity and geographical distribution (Shinozaki and Yamaguchi-Shinozaki 2000). *GhNAC4* is significantly upregulated by cold stress and drought and its promoter region showed many motifs like MYB2CONSENSUSAT, C-REPEAT/DRE, DRECRTCOREAT, CSD, and CBFHV TF binding sites required for drought and cold responsiveness. Two copies of MYB2CONSENSUSAT motifs are found in the *GhNAC4* promoter, which is the binding site for AtMYB2 TF, that is required for drought inducibility of *rd22* gene by binding to its promoter region (Abe 2003). Cold responsive motif, C-repeat (CRT) is responsible for the regulation of many cold-inducible genes in an ABA-independent manner. It is also involved in dehydration responsiveness (Stockinger et al. 1997). An AP2 domain-containing TF, CBF/DREB binds to this motif (Liu et al. 1998). Three copies of the DRECRTCORE motif are found in the *GhNAC4* promoter. CBFHV motif is a binding site of an AP2 domain containing cold-inducible TF, HvCBF1 characterized in barley (Xue 2002). Three copies of CBFHV motif are found in the *GhNAC4* promoter. Eight copies of binding site for cold shock domain (CSD) proteins are also found in the *GhNAC4* promoter that is highly upregulated during low-temperature stress. Apart from being important during cold adaptation, they are also known to play roles in plant development (Chaikam and Karlson 2008).

Heat stress damages the components of the photosynthetic apparatus and leads to oxidative stress, thereby affecting plant growth and productivity (Kotak et al. 2007). High temperature and oxidative stress induced the *GhNAC4* transcripts by several folds. Heat stress proteins (HSP) are molecular chaperones that are essential for restoring homeostasis during heat stress, and the transcriptional regulation of HSPs is controlled by heat stress transcription factors (HSF, Al-Whaibi 2011). They bind to the heat stress responsive element (HSE) and modulate transcription during heat stress (Baniwal et al. 2004). Four copies of HSF binding site are present in the *GhNAC4* promoter. Binding sites for C2H2 TF were also identified in the *GhNAC4* promoter region. C2H2 TFs such as ZAT7, ZAT10, and ZPT2 are known to be important in regulating responses to abiotic and biotic stress tolerance (Kiełbowicz-Matuk 2012).

Plants defend themselves against pathogen attacks by developing strong physical barriers in their cell walls and a complex signalling network that involves inducible defence mechanisms (Yang et al. 1997). *GhNAC4* promoter contains several binding sites of TFs like WRKY, MYB, BELL and also many copies of motifs such as WBOXNTCHN48, BOXLCROREDCPAL, HSRENTHSR203J and GT1GMSCAM4 known to be essential for elicitor-induced activation of defence genes (Eulgem 2005).

Elicitor induced WRKY TF is involved in the transcription of defence genes like chalcone synthase (*CHN48*) in tobacco by binding to the WBOXNTCHN48 motif (Yamamoto et al. 2004). DcMYB1 TF binds to the BOXLCROREDCPAL motif in the promoter of phenylalanine ammonia-lyase (*PAL*) gene in carrot and is essential for induction by elicitor treatment or UV-B irradiation (Maeda et al. 2005). *HSR203J* gene is rapidly and specifically upregulated during a hypersensitive response and HSRENTHSR203J (HSR203 responsive element) motif is responsible for the induction of HSR203J during incompatible plant-pathogen interactions (Pontier et al. 2001). Soybean calmodulin (SCaM-4) is rapidly induced by pathogen attack and salt stress. This is mediated by the binding of GT-1 TF to the promoter at the GT1GMSCAM4 motif (Park et al. 2004). Six such motifs are also present in the *GhNAC4* promoter. OsBIHD1 TF is a BELL homeodomain TF that is known to induce resistance in rice in response to *Magnaporthe grisea* (Luo et al. 2005). Six copies of OsBIHD1 binding sites been observed in the *GhNAC4* promoter. Mechanical wounding induced the expression of *GhNAC4* by several folds and we observed intense GUS staining in the wounded area in the transgenic tobacco plants that expressed the PRO*_GhNAC4_*:GUS fusion. *GhNAC4* promoter exhibited four copies of WBOXNTERF3 motifs. WRKY TF binds to the WBOXNTERF3 motif in the promoter of *ERF3* gene of tobacco and causes its rapid activation upon wounding (Nishiuchi et al. 2004). Thus, these motifs may modulate the expression of *GhNAC4* gene in response to various abiotic and biotic stresses.

The presence of ABA-dependent and independent abiotic stress-related motifs and also elicitor-induced motifs in the *GhNAC4* promoter suggests that there is a cross-talk between various stress signalling pathways and *GhNAC4* may act as a node for the integration of these pathways. This may be modulated by the interactions of different motifs present in the *GhNAC4* promoter.

### Tissue and organ-specific expression of the *GhNAC4* promoter

Stress responses occur primarily at the level of transcription leading to the regulation of spatial and temporal expression of stress-induced genes (Geng et al. 2013). Spatial and temporal patterns under the control of *GhNAC4* promoter were monitored in various tissues by the detection of GUS activity in transgenic tobacco plants. In the present investigation, we have observed the expression of *GhNAC4* at the seedling stage, in the emerging radicle, plumule and cotyledons. *GhNAC4* expression was also observed in the vascular bundles of the stem, leaf, main and lateral roots, in the later stages of plant growth. It was also observed in the guard cells and meristems. During flowering, sepals, petal edges, and pollens showed *GhNAC4* expression. The occurrence of very strong vasculature specific activities for *GhNAC4* promoter could be corroborated by the presence of several motifs responsible for these tissue-specific expression patterns. Several copies of SEF1MOTIF, SEF3MOTIFGM, and SEF4MOTIFGM7S motifs were observed in the *GhNAC4* promoter. These are developing embryo and seed-specific motifs found upstream of soybean seed storage β-conglycinin gene (Lessard et al. 1991). Binding sites for three-amino-acid-loop-extension (TALE) TFs that are known to control the formation and maintenance of meristem are observed in the promoter of *GhNAC4* gene. These homeo-proteins provide a gene regulatory link between hormonal stimuli and development of shoot apical meristem (Hamant and Pautot 2010).

OSE1ROOTNODULE and OSE2ROOTNODULE motifs are root specific elements found in the promoter of *Vicia faba* leghaemoglobin gene and were shown to be important for nodule formation by arbuscular mycorrhiza (Fehlberg et al. 2005). The ROOTMOTIFTAPOX1 element is found in the promoter of the *rolD* gene of *Agrobacterium rhizogenes,* which has a distinct expression pattern in the root elongation zone and vascular bundle (Elmayan and Tepfer 1995). The CACTFTPPCA1 motif is a key component of mesophyll expression module 1 (MEM1) in the promoter of *Flaveria trinervia* C4 system based phosphoenolpyruvate carboxylase gene and is sufficient for high mesophyll-specific expression (Gowik et al. 2004). Twenty-two copies of CACTFTPPCA1 motif are observed in the *GhNAC4* promoter region. The TAAAGSTKST1 is a guard cell-specific motif found upstream of K ^+^ influx channel gene (*KST1*) in potato and is bound by StDof1 TF (Plesch et al. 2001). Five copies of TAAAGSTKST1 motif are also observed in the *GhNAC4* promoter region.

Binding sites for quite a few other TF known to play roles in organogenesis and tissue-specific expressions were predicted such as MYBST1, DOFCOREZM, SBP TF, TCR TF, AT HOOK TF, and WOX TF. *GhNAC4* has 26 copies of DOFCOREZM motif, which is a binding site of DOF TFs that are plant specific and have a unique single zinc finger DNA binding domain. A key role of DOF transcription factors in the formation and functioning of plant vascular bundles is emerging (Le Hir and Bellini 2013). They regulate directly or indirectly the processes associated with the establishment and maintenance of the vascular system (Le Hir and Bellini 2013). HMG-I/Y TFs have an AT-hook region, which is a short conserved peptide that binds to AT-rich tracts of DNA in the minor groove and cause DNA bending (Reeves 2000). These TFs play an important role in chromatin structure and regulate gene expression by acting as TF co-factors (Reeves and Beckerbauer 2001). HMG-I/Y TFs are expressed in high levels in tissues showing rapid cell division. And, they are also known to play an important role in development and defence responses (Klosterman and Hadwiger 2002).

Squamosa promoter binding protein (SBP) TFs are plant-specific TFs that are known to play a role both in vegetative phase change and flower development pathway and their role in stress response is also emerging (Hou et al. 2013; Klein et al. 1996). Wuschel-related homeobox (WOX) TFs play important roles in cell fate determination during all stages of plant development and shows very specific spatial and temporal expression (Hedman et al. 2013). Various motifs such as CARCGW8GAT, TGTCACACMCUCUMISIN and binding sites for MYBPZM and TCR TFs known to be important for flower and fruit development were found. CARCGW8GAT is the binding site of MADS-box TF, AGAMOUS-like 15 (AGL15). AGl15 accumulates during embryo development and regulates somatic embryogenesis (Tang and Perry 2003). TGTCACACMCUCUMISIN is a fruit-specific motif found upstream of a subtilisin-like serine protease, a cucumisin gene in melon (Yamagata et al. 2002). MYBPZM is an MYB homolog, P, transcription factor from maize that binds to CCT/AACC sequence and it controls red pigmentation in the floral organs by regulating the flavonoid biosynthetic pathway (Grotewold et al. 1994). Two pollen-specific motifs are also found in the *GhNAC4* promoter, POLLEN1LELAT52, and GTGANTG10. POLLEN1LELAT52 is one of the two co-dependent motifs found upstream of tomato *LAT52* gene that is essential for pollen development (Bate and Twell 1998). The GTGANTG10 motif is important for the pollen-specific expression of tobacco pectate lyases, *g10* gene (Rogers et al. 2001). GhNAC4 has a unique tissue-specific expression pattern suggesting that this gene may be essential for plant growth and development.

### Sugar-responsive motifs in *GhNAC4* promoter

In plants, soluble sugars not only function as a nutrient source but also act as signals regulating various growth and development pathways (Rolland et al. 2006). There is a cross-talk between the environmental stress response and the sugar signalling pathways to modulate plant metabolic responses by differential regulation of many genes (Ho et al. 2001). Nutritional stress and sugar starvation responses may have an overlap with the response of a plant to environmental stresses (Couée et al. 2006). Exogenous application of glucose helped alleviate of the negative effects of salt stress in wheat seedlings by maintaining the ion homeostasis and activation of the antioxidant system (Hu et al. 2012). *GhNAC4* promoter exhibits 16 copies of CGACGOSAMY, two copies of WBOXHVISO1 and one copy of SREATMSD motifs. CGACGOSAMY3 motif acts as a coupling element for the G-box motif in the promoter of sugar starvation regulated rice amylase 3D (*OsAMY3D*) gene and plays a role in sugar responsiveness coupled to environmental stress regulation (Hwang et al. 1998). SUGAR SIGNALLING IN BARLEY 2 (*SUSIBA2*) is a sugar-inducible WRKY TF, which binds to both WBOX (*WBOXHVISO1*) and SURE (Sugar responsive element) in the barley isoamylase1 (*iso1*) promoter and is involved in the regulation of starch synthesis (Sun et al. 2003). SREATMSD is a sugar-repressive element, found upstream of genes down-regulated during auxiliary bud growth after main stem decapitation. SREATMSD is also involved in feed-back sugar repression of gene expression (Tatematsu et al. 2005). Sugar-dependent repression of gene expression is an important regulatory mechanism for adjusting the plant to the changes in the availability of carbon source and maintaining energy homeostasis between the source and sink tissues (Yu et al. 2015).

### Light-responsive elements in *GhNAC4* promoter

Plants responses to light are complex, and most of these responses such as development and stress signalling require the regulation in the expression of various nuclear and plastid-encoded genes and are mediated by photoreceptors (Kong and Okajima 2016). More than one photoreceptors may act on light-regulated genes thereby allowing tight control of their expression upon light stimulation (Kami et al. 2010). It has been hypothesised that light-responsive elements (LRE) have complex structures and consist of aggregates of connected binding sites for different TFs (Terzaghi and Cashmore 1995). Combinatorial interaction of different LREs in the promoter sequence such as GATA motif, ASL box, BOXII, IBOX, GT1 element, SORLIPs, TBOX, AE-BOX, and ATC motif, is required to confer light inducibility (Argüello-Astorga and Herrera-Estrella 1998). The combinatorial pairing of tetrameric repeats of GATA and GT1 motif to a minimal promoter responded to a broad spectrum of light as compared to multimeric repeats of these motifs alone (Chattopadhyay et al. 1998). GATA motif plays a role in light responsiveness and tissue specificity and is involved in the light-dependent development of phloem tissue (Yin et al. 1997). Vascular localisation of *GhNAC4* (Fig. 6) could be partially attributed to the presence of 42 copies of the GATA motif in its promoter region. Several copies of GT1CONSENSUS motif were found in the *GhNAC4* promoter sequence. GT1CONSENSUS sequence (GRWAAW) was initially identified as Box II motif and is the binding site of *GT-1* TF. It is involved in light activation or dark repression and tissue-specificity (Hiratsuka et al. 1994). SORLIPs (sequence over-represented in light-induced promoters) are light-responsive elements found in the promoters of genes that are involved in the phytochrome A receptor pathway (Hudson and Quail 2003). SORLIP1AT and SORLIP2AT have been noted in the *GhNAC4* promoter sequence. TBOXATGAPB motif is found upstream of Arabidopsis *GAPDH* subunit B gene and serves as a positive modulator of light inducibility (Chan et al. 2001). This suggests that the tissue specificity of GhNAC4 to a certain extent is dependent also on light inducibility.

Abiotic stress impedes plant growth and development which would compel the regulation of photosynthesis and carbohydrate partitioning. *GhNAC4* showed the presence of sugar and light regulatory motifs and is also up-regulated by abiotic stresses. This indicates that GhNAC4 might act in integrating stress responses with plant growth regulation.

Thus, the presence of key motifs related to various stresses in the promoter of *GhNAC4* indicates their putative function in the response of a cotton plant to various environmental stresses. The transcript abundance of *GhNAC4* was altered by salinity, osmotic stress, oxidative stress, wounding, low and high temperatures and drought, as well as, exogenous application of various phytohormones like ABA, BAP, MeJA, GA and ethylene implying that *GhNAC4* may participate in the response of the cotton plant’s to environmental stress. This work would provide a foundation for a comprehensive functional investigation of NAC TFs in the future.

## Conclusion

In the present study, we showed that *GhNAC4* expression is responsive to phytohormones like ABA, JA, CK, and auxin and is up-regulated under external stimuli like drought, oxidative stress, osmotic stress, salinity, and cold. We also demonstrated that *GhNAC4* promoter is a vascular-specific promoter by histochemical assay and thin sectioning. Further analysis of GUS activity showed that GhNAC4 promoter could be induced by multiple plant hormones and environmental stresses. Bioinformatic analysis predicted the presence of various *cis*-regulatory elements associated with tissue specificity, phytohormone responsiveness, stress inducibility, light inducibility and sugar responsiveness suggesting that GhNAC4 TF may be a common regulator of the molecular mechanism controlling plant development and stress responses. This work provides a solid foundation for the use of *GhNAC4* promoter in biotechnological approaches as a vasculature-specific and multiple stress-inducible promoter. We are currently investigating the effects of GhNAC4 expression in transgenic plants and how GHNAC4 integrates stress responses with plant developmental processes.

## Abbreviations

4-MU: 4-Methylumbelliferone
ABA: Abscisic acid
BAP: 6-Benzyl aminopurine
GA: Gibberellic acid
GUS: β-Glucuronidase
IAA: Indole-3-acetic acid
MeJA: Methyl Jasmonic acid
MUG: 4-Methylumbelliferyl-β-Dglucuronide
MV: Methyl viologen
NAC: NAM ATAF CUC
PEG: Polyethylene glycol
SA: Salicyclic acid
TF: Transcription factor

## Author contribution statement

VST, GP and PBK conceived and designed the experiments. VST, SM and PB performed the experiments. VST analysed the data. VST and PBK wrote the manuscript. All authors read and approved the manuscript.

## Acknowledgements

TSV would like to acknowledge Council of Scientific and Industrial Research (CSIR) for research fellowship. Also, Vikas Kumar Jain and Manimaran Panneer Selvam for critical reading of the manuscript. The authors are grateful to the Head, Department of Plant Sciences, University of Hyderabad for the facilities under various umbrella programs like DST-FIST and UGC-SAP-DRS.

## Conflct of interest

The authors declare they have no conflict of interest.

